# Simple Behavioral Analysis (SimBA) – an open source toolkit for computer classification of complex social behaviors in experimental animals

**DOI:** 10.1101/2020.04.19.049452

**Authors:** Simon RO Nilsson, Nastacia L. Goodwin, Jia Jie Choong, Sophia Hwang, Hayden R Wright, Zane C Norville, Xiaoyu Tong, Dayu Lin, Brandon S. Bentzley, Neir Eshel, Ryan J McLaughlin, Sam A. Golden

## Abstract

Aberrant social behavior is a core feature of many neuropsychiatric disorders, yet the study of complex social behavior in freely moving rodents is relatively infrequently incorporated into preclinical models. This likely contributes to limited translational impact. A major bottleneck for the adoption of socially complex, ethology-rich, preclinical procedures are the technical limitations for consistently annotating detailed behavioral repertoires of rodent social behavior. Manual annotation is subjective, prone to observer drift, and extremely time-intensive. Commercial approaches are expensive and inferior to manual annotation. Open-source alternatives often require significant investments in specialized hardware and significant computational and programming knowledge. By combining recent computational advances in convolutional neural networks and pose-estimation with further machine learning analysis, complex rodent social behavior is primed for inclusion under the umbrella of computational neuroethology.

Here we present an open-source package with graphical interface and workflow (Simple Behavioral Analysis, SimBA) that uses pose-estimation to create supervised machine learning predictive classifiers of rodent social behavior, with millisecond resolution and accuracies that can out-perform human observers. SimBA does not require specialized video acquisition hardware nor extensive computational background. Standard descriptive statistical analysis, along with graphical region of interest annotation, are provided in addition to predictive classifier generation. To increase ease-of-use for behavioural neuroscientists, we designed SimBA with accessible menus for pre-processing videos, annotating behavioural training datasets, selecting advanced machine learning options, robust classifier validation functions and flexible visualizations tools. This allows for predictive classifier transparency, explainability and tunability prior to, and during, experimental use. We demonstrate that this approach is flexible and robust in both mice and rats by classifying social behaviors that are commonly central to the study of brain function and social motivation. Finally, we provide a library of poseestimation weights and behavioral predictive classifiers for resident-intruder behaviors in mice and rats. All code and data, together with detailed tutorials and documentation, are available on the <u>SimBA GitHub repository</u>.

**Graphical abstract:** **SimBA graphical interface (GUI) for creating supervised machine learning classifiers of rodent social behavior.**

*(a) <u>Pre-process videos</u>*. SimBA supports common video pre-processing functions (e.g., cropping, clipping, sampling, format conversion, etc.) that can be performed either on single videos, or as a batch.

*(b) <u>Managing poseestimation data and creating classification projects</u>*. Pose-estimation tracking projects in DeepLabCut and DeepPoseKit can be either imported or created and managed within the SimBA graphical user interface, and the tracking results are imported into SimBA classification projects.

SimBA also supports user-drawn region-of-interests (ROIs) for descriptive statistics of animal movements, or as features in machine learning classification projects.

*(c) <u>Create classifiers, perform classifications, and analyze classification data</u>*. SimBA has graphical tools for correcting poseestimation tracking inaccuracies when multiple subjects are within a single frame, annotating behavioral events from videos, and optimizing machine learning hyperparameters and discrimination thresholds. A number of validation checkpoints and logs are included for increased classifier explainability and tunability prior to, and during, experimental use. Both detailed and summary data are provided at the end of classifier analysis. SimBA accepts behavioral annotations generated elsewhere (such as through JWatcher) that can be imported into SimBA classification projects.

*(d) <u>Visualize classification results</u>*. SimBA has several options for visualizing machine learning classifications, animal movements and ROI data, and analyzing the durations and frequencies of classified behaviors.

See the <u>SimBA GitHub repository</u> for a comprehensive documentation and user tutorials.

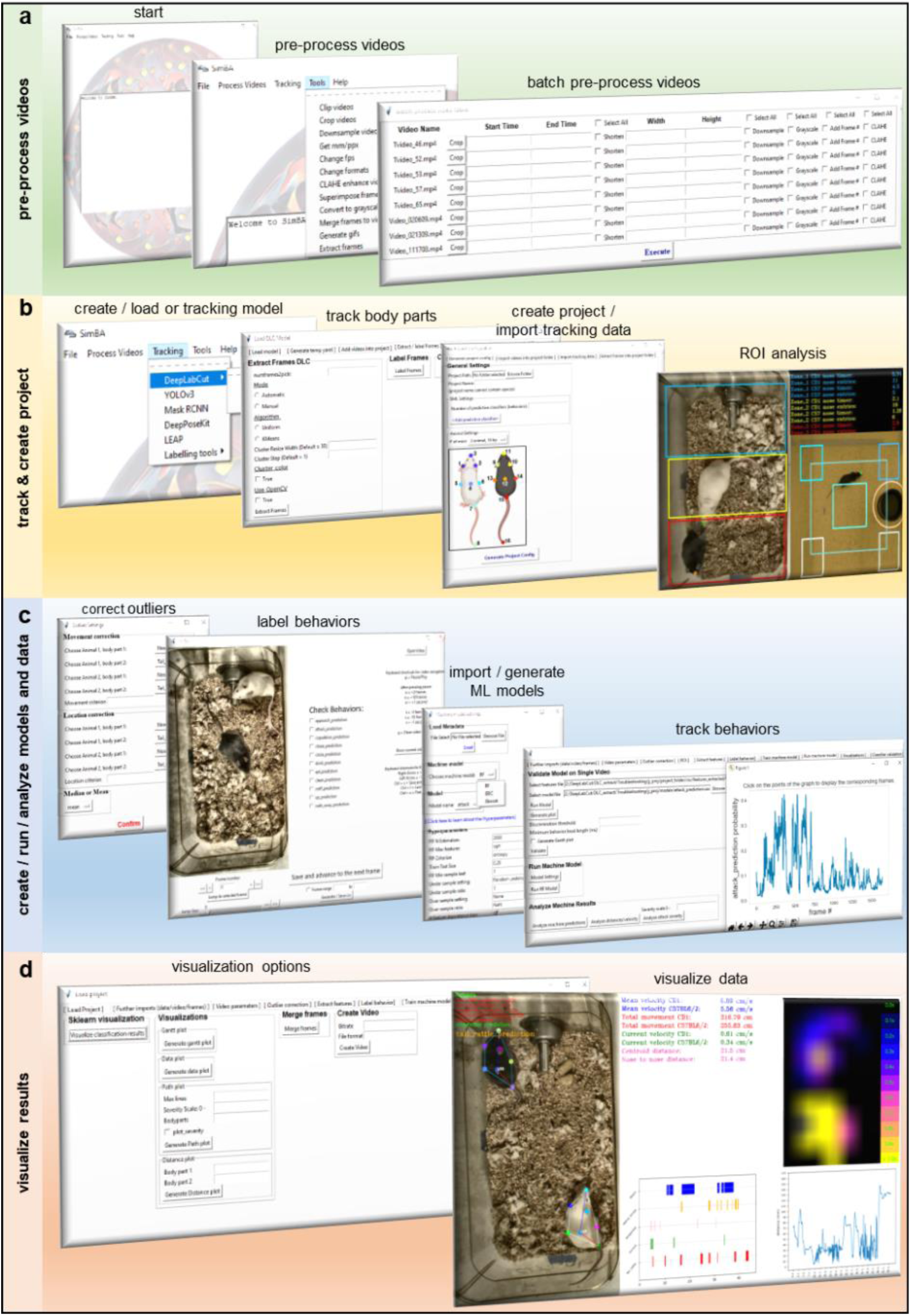

## Introduction

Maladaptive social behaviours are theorised as both a cause and consequence of many neuropsychiatric disorders including depression^1,2^, substance abuse^3^ and PTSD^4–7^. Such social impairments can result from compromised reward evaluations where the motivation for maintaining healthy relationships are persistently diminished^8^. As such, the treatment of maladaptive social behaviour presents an important strategy for improving mental health^9–12^. However, despite the established role of social behaviour in psychiatric disorders and treatments, it remains unclear how social context interact with biological functions to produce protracted and chronic sequalae^10^. These insights depend on experimental animal model systems where detailed behavioral ethograms can be combined with precise manipulations of brain functions to reveal biological causes^12–14^.

Unfortunately, the preclinical study of complex social behavior is impeded by the difficulties associated with reliably scoring complex social interactions. Analyses are typically performed by a trained individual, or preferably several trained individuals for appropriate inter-rater-reliability validation, observing social behaviors and manually scoring previously defined events and their durations. The approach is arduous, non-standardized and susceptible to confounds produced by observer drift, long analysis times, and poor inter-rater-reliability. These confounds often prevent the detailed study of complex social repertoires in larger datasets, and notably provide far lower temporal resolution than most modern methodologies such a in vivo electrophysiological recording, fiber photometry recording and single-cell calcium endomicroscopy^15–18^. These important omissions may contribute to our limited biological understanding of maladaptive social behavior and their neurobiological underpinnings.

To overcome this, a range of elegant open-source tools (Table 1) that use various forms of computer vision with synchronized RFID-tracking data and/or depth camera or multi-camera 3D systems have been developed for automated and precise classifications of animal behaviour^19–24^. Such methods can permit online classifications in semi-natural environments and are a foundation for impending closed-loop monitoring and forecasting systems^25^ in behavioural neuroscience, but do require significant investment in specialized hardware and a working knowledge of computer science approaches. Parallel advances in animal tracking have produced accessible open-source pose-estimation tools for accurate tracking of experimenter-defined body-parts in noisy and variable environments^26–29^ (Table 1). These open-source initiatives have proven to be both far less expensive to execute, and provide better animal tracking outcomes, than currently available commercial products^30^. The collective developments promise decreased workload and reduced experimenter interference while introducing sampling frequencies and scoring accuracies that are compatible with other modern neuroscience techniques. Automated classification techniques may also reduce confounds and increase replicability through computer models based on standardized cross-laboratory definitions of behavioral repertoires, made available in libraries for the scientific community. Together, these approaches may solidify the study of rodent social behavioral under the umbrella of the recently-reinvigorated field of computational neuroethology^31–36^ and increase our understanding of naturalistic unrestrained behavior within the context of neural function.

**Table 1.** Open-source computational packages for animal tracking and/or behavioral classification (not comprehensive)

| Name | Validated Species | Multiple animals? | Main output | Behavioral classifiers validated in original publication | Mechanism |
| --- | --- | --- | --- | --- | --- |
| **Ctrax**<sup>87</sup> | Drosophila | Yes | Behavior | Walk, stop, sharp turn, crabwalk, backup, touch, chase | Computer vision |
| **CADABRA**<sup>88</sup> | Drosophila | Yes | Behavior | Lunging, tussling, wing threat/extension, circling, chasing, copulation | Supervised |
| **Autotyping**<sup>89</sup> | Mice | No | Behavior | Home-cage behaviors: drink, eat, groom, hang, micromovement, rear, rest, walk | Supervised |
| **JAABA**<sup>24</sup> | Drosophila, mice | Yes | Behavior | Flies: walk, stop, crabwalk, backup, touch, chase, jump, copulation, wing flick/grooming/extension, righting, center pivot, tail pivot. Mice: follow, walk. | Supervised |
| **MiceProfiler**<sup>90</sup> | Mice | Yes, if no obstruction | Behavior | Head-head contact, anogenital contact, side by side, approach, leave, follow, chase | Physics engine |
| **Unnamed**<sup>*91</sup> | Mice | Yes | Behavior | Approach, walk away, circle, chase, attack, copulation, drink, eat, clean, sniff, rear | Supervised |
| **Unnamed**<sup>*92</sup> | Rats | Yes | Behavior | Rearing, head-head and head – hip contact, approach, leave, follow, mount, intromission, ejaculation | Physics engine |
| **MotionMapper**<sup>93</sup> | Flies | Yes | Behavior | Grooming: wing, leg, abdomen, head. Wing waggle, running | Unsupervised |
| **Unnamed**<sup>*94</sup> | Mice | Yes, if distinct | Behavior | Attack, close investigation, mounting | Supervised |
| **MoSeq**<sup>*95</sup> | Mice | No | Behavior | Dart, micromovement, pause, rear, walk | Unsupervised |
| **Unnamed**<sup>96</sup> | Mice | Yes | Behavior | Direct contact, nose to nose, anogenital sniffing, following, mating, self-grooming | Unsupervised |
| **DuoMouse**<sup>97</sup> | Mice | Yes | Behavior | Sniffing, following, indifferent | Unsupervised |
| **Stytra**<sup>98</sup> | Zebrafish larvae | No | Closed loop tracking & stimulus admin. (real time) |  | Computer vision |
| **ToxID / ToxTrac**<sup>99,100</sup> | Fish, ants, mice | Yes | Individual tracking in groups (real time) |  | Computer vision |
| **MoST**<sup>101</sup> | Mice | Yes | Individual tracking in groups |  | Computer vision |
| **idtracker**<sup>75,76</sup> | Drosophila, fish, ants, mice | Yes | Individual tracking in groups |  | Unsupervised |
| **LocoMouse**<sup>*102</sup> | Mice | No | Kinematics |  | Supervised |
| **DeepLabCut**<sup>27</sup> | Drosophila, mice, horses, humans, fish | Expected soon | Pose |  | Supervised |
| **LEAP**<sup>28</sup> | Drosophila, mice | sLEAP expected soon | Pose |  | Supervised |
| **DeepFly3d**<sup>* 103</sup> | Drosophila, humans | No | Pose |  | Unsupervised |
| **DeepPoseKit**<sup>70</sup> | Drosophila, locusts, zebras | Yes, if distinct | Pose (real time) |  | Supervised |
| **Sensory Orientation Software**<sup>104</sup> | Drosophila, mice, fish | No | Posture & trajectory (real time) |  | Computer vision |
| **MOTR**<sup>105</sup> | Mice | Yes, if distinct | Trajectory |  | Computer vision |
| **Mouse Behavior Tracker**<sup>106</sup> | Mice | No | Trajectory |  | Computer vision |

Asterisk denotes requirement of multiple cameras or special equipment

However - despite these developments - manual experimenter annotations persist as the conventional approach for studying social behavior in experimental animals. We have identified a number of interrelated obstacles that prevent widespread implementation of machine learning techniques for behavioral neuroscience, and we are presenting this perspective in an upcoming in-depth review^37^. Now briefly, and most importantly, these include (i) the moderate to advanced computational/engineering skills and significant time investments that are necessary to implement most machine learning approaches, and (ii) the “black-box” nature and lack of explainability^38,39^ and validation that current approaches are often paired with. Put simply, the learning curves for novice users are steep and often dependent on skill-sets not commonly taught within behavioral neuroscience curricula, and the interpretation of their outcomes can be ethereal.

Here we present an open-source method and graphical user-interface (GUI) called SimBA (**Sim**ple **B**ehavioral **A**nalysis; Fig. 1) that is used to generate supervised decision ensembles of complex social behaviors from basic video recordings of mouse and rat dyadic encounters. SimBA uses features generated from pose-estimation tracking data together with experimenter-made annotations to generate random forests algorithms that accurately classify behavioral patterns in experimental videos. We package SimBA with a range of accessible tools for processing video data and advanced functions for validating and evaluating predictive classifiers and visualizing machine classifications. SimBA has been developed for analysing complex social interactions but has been adopted for a variety of non-social behavioral protocols at other labs and institutes. We show that the method is flexible by accurately predicting an extensive set of behaviours relevant for studying the neural mechanisms underlying social motivations in mice and rats. All methods, code, and data together with detailed tutorials are available on the <u>SimBA GitHub repository</u>.

**Figure 1.**
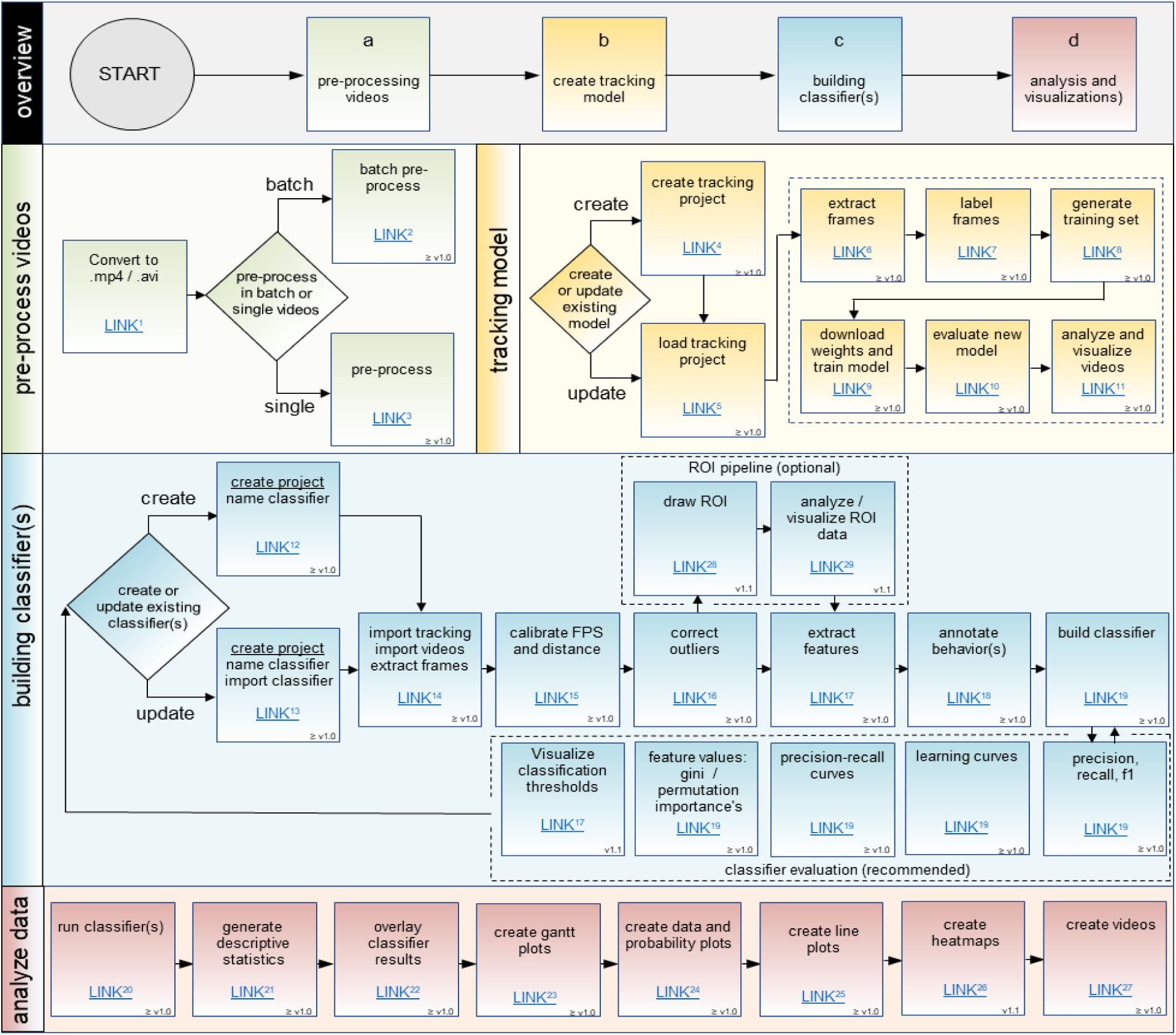
Flowchart depicting the SimBA workflow for creating supervised machine learning classifiers. An active-link version of this figure is found on the <u>SimBA GitHub repository</u>. *(a) Pre-process videos*. SimBA provided built-in pre-processing options (trim, crop, enhance, down-sample) that can be applied to either single or multiple videos prior to pose-estimation body-part tracking. Videos are CLAHE (Contrast Limited Adaptive Histogram Equalization) enhanced if recorded in greyscale and/or deficient resolution. *(b) Animal tracking*. The pre-processed videos are analyzed using the appropriate DeepLabCut/DeepPoseKit pose-estimation model. Pretrained pose-estimation model weights for a variety of experimental protocols are available to download from the <u>SimBA OSF repository</u>. *(c) Building classifiers*. Gross inaccuracies in pose-estimation tracking are corrected, and machine learning features (e.g., movements, distances etc.) are calculated from the corrected tracking data. The videos are annotated for the behaviors of interest using the SimBA event-logger, and the annotated events are concatenated with the corrected pose-estimation tracking data. Random forest hyperparameters and discrimination thresholds are tuned, and classifier performances are evaluated. Our random forest classifiers for resident-intruder protocols can be imported into SimBA projects and are available to download from the <u>SimBA OSF repository</u>. *(d) Analysis and visualizations*. The patterns of the classified behaviors, and critical machine learning features (velocities, total movements etc.), are analyzed and visualized. See <u>SimBA on YouTube</u> for visualization examples.

## Methods

### SimBA workflow overview

A flowchart depicting the workflow of SimBA is shown in Fig. 1, with links to access online tutorials for the specific SimBA modules. An active-link version of Fig. 1 is found on the <u>SimBA GitHub repository</u>.

#### Hardware requirements

The recommended system requirements and detailed installation instructions can be found on the <u>SimBA GitHub repository</u>. Briefly, the minimum specifications are an i7 CPU, 16GB RAM, and SSD hard drive. The recommended specifications are an i9 CPU, 32GB RAM, an SSD hard drive, and NVIDEA RTX2080Ti GPU. Animal pose-estimation tracking (i.e., DeepLabCut^27^ / DeepPoseKit^26^), component of the SimBA pipeline, requires GPU support. Importantly, SimBA can accept imported pose-estimation data that has been previously generated or generated on a separate system. The visualization steps of SimBA rely on basic functions in the OpenCV and FFmpeg libraries, with are free software suits for video editing and computer vision, and benefit from higher-end CPUs and SSDs.

#### Installation options

Two supported versions of SimBA (Fig. 2) are available to download from the <u>SimBA GitHub repository</u>: *SimBAxTF* and *SimBA*. The *SimBAxTF* version requires TensorFlow^40^, local GPU-support, and has integrated graphical menus for creating convolutional neural networks through the DeepLabCut^27^ and DeepPoseKit^26^ packages. Conversely, the self-contained SimBA version does not require TensorFlow, or local GPU-support, and accepts imported pose-estimation tracking data generated separately within the DeepLabCut or DeepPoseKit notebooks and interfaces.

**Figure 2.**
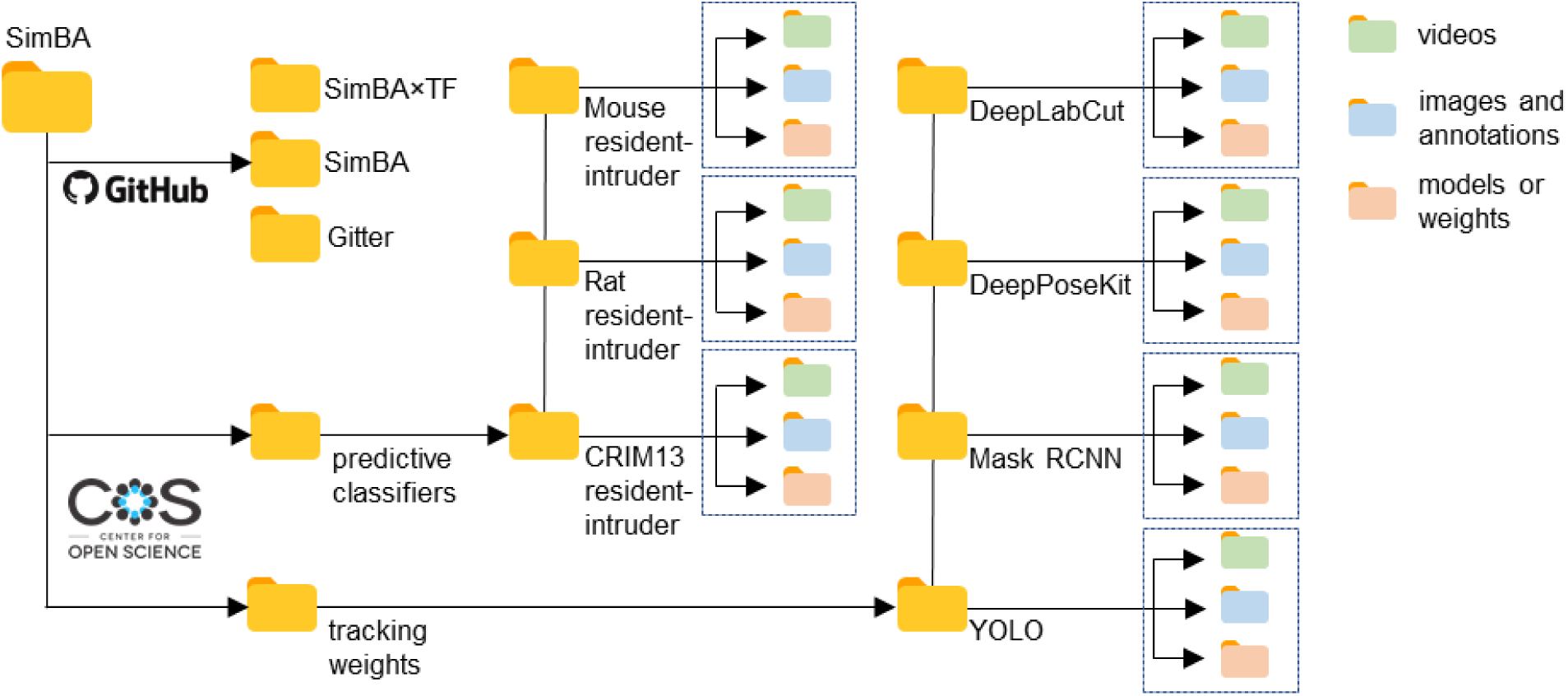
The SimBA open-source code-base and folder structure of the open-access datasets for creating social behavior predictive classifiers. Two supported versions of SimBA are available to download from the <u>SimBA GitHub repository</u>; SimBA with (SimBA×TF) and without (SimBA) integrated TensorFlow-support for convolutional neural networks and pose-estimation animal tracking through DeepLabCut and DeepPoseKit. Support is available through the <u>SimBA *Gitter*</u> instant messaging service and chat room. Go to the <u>SimBA Open Science Framework (OSF) repository</u> to download the annotations, videos, random forest models, and tracking weights within predictive classification projects. The <u>SimBA OSF repository</u> also contains image annotations and network weights for tracking white and black coat-colored animals using alternative neural network architectures (e.g., mask RCNN and YOLO).

#### Software

SimBA is written for Windows compatibility using only open-source software. The code was written in python3^41^. The GUI was written in tkinter^42^. The GUI uses DeepLabCut^27^ or DeepPoseKit^26^ for body-part labeling and body-part pose-estimation. The SimBA code for generating machine learning behavior classifiers and visualizations rely primarily on scikit-learn^43^, OpenCV^44^, FFmpeg^45^, and imblearn^46^. A list of SimBA package dependencies and detailed installation instructions can be found on the <u>SimBA GitHub repository</u>.

#### Datasets

We used SimBA to create social predictive classifiers from three experimental protocols (Table 2): mouse resident-intruder, rat resident-intruder, and the <u>Caltech Resident-Intruder Mouse dataset (CRIM13)</u>^19,47^.

**Table 2.** Predictive classifier datasets

|  | classifier | # annotated frames | # annotated videos | annotations: % present | annotations: present (hh:mm:ss) | annotations: absent (hh:mm:ss) | under sample ratio | test set: frames present | test set: frames absent | optimal threshold |
| --- | --- | --- | --- | --- | --- | --- | --- | --- | --- | --- |
| mouse resident-intruder | allogroom normal | 334680 | 26 | 1.49 | 00:02:46 | 03:03:10 | 19 | 983 | 65953 | 0.48 |
|  | allogroom vigorous | 334680 | 26 | 0.73 | 00:01:21 | 03:04:35 | 0 | 494 | 66442 | 0.41 |
|  | attack | 203841 | 37 | 4.93 | 00:05:35 | 01:47:40 | 9 | 1920 | 38849 | 0.56 |
|  | lateral threat | 168738 | 21 | 1.49 | 00:01:24 | 01:32:21 | 18 | 503 | 33245 | 0.52 |
|  | mounting | 470294 | 31 | 2.88 | 00:07:31 | 04:13:45 | 4 | 2639 | 91420 | 0.72 |
|  | pursuit | 103538 | 33 | 2.02 | 00:01:10 | 00:56:22 | 20 | 419 | 20289 | 0.43 |
|  | upright submissive | 272304 | 20 | 1.13 | 00:01:43 | 02:29:34 | 16 | 577 | 53884 | 0.54 |
|  | flee | 395852 | 22 | 1.08 | 00:02:23 | 03:37:33 | 28 | 877 | 78294 | 0.60 |
|  | scramble | 323852 | 18 | 2.16 | 00:03:53 | 02:56:02 | 3.9 | 1347 | 63424 | 0.79 |
|  | anogenital sniffing | 103538 | 33 | 6.57 | 00:03:47 | 00:53:45 | 8 | 1268 | 19440 | 0.55 |
| rat resident-intruder | attack | 136830 | 12 | 16.40 | 00:12:28 | 01:03:33 | 0 | 4437 | 32119 | 0.55 |
|  | lateral threat | 136830 | 12 | 9.09 | 00:06:55 | 01:09:06 | 0 | 9943 | 26613 | 0.55 |
|  | anogenital sniffing | 136830 | 12 | 7.50 | 00:05:42 | 01:10:19 | 7.5 | 2095 | 22713 | 0.45 |
|  | submission | 136830 | 12 | 11.27 | 00:08:34 | 01:07:27 | 5 | 3074 | 33482 | 0.59 |
|  | boxing | 136830 | 12 | 4.50 | 00:03:25 | 01:12:36 | 0 | 1306 | 23502 | 0.47 |
|  | approach | 136830 | 12 | 4.23 | 00:03:13 | 01:12:48 | 8.5 | 1053 | 23755 | 0.50 |
|  | avoidance | 136830 | 12 | 2.48 | 00:01:53 | 01:14:08 | 10 | 686 | 24122 | 0.48 |
| CRIM13 resident-intruder | approach | 757350 | 64 | 4.13 | 00:20:51 | 18:01:39 | 12 | 6190 | 145280 | 0.44 |
|  | attack | 757350 | 64 | 4.10 | 00:20:42 | 18:01:48 | 16 | 6266 | 145204 | 0.40 |
|  | chase | 757350 | 64 | 0.49 | 00:02:28 | 18:20:02 | 46 | 769 | 150701 | 0.58 |
|  | circle | 757350 | 64 | 1.32 | 00:06:40 | 18:15:50 | 22 | 1982 | 149488 | 0.54 |
|  | clean | 757350 | 64 | 7.99 | 00:40:20 | 17:42:10 | 8 | 12095 | 139375 | 0.53 |
|  | copulation | 757350 | 64 | 4.62 | 00:23:20 | 17:59:10 | 12 | 7080 | 144390 | 0.50 |
|  | drink | 757350 | 64 | 0.36 | 00:01:49 | 18:20:41 | 10 | 570 | 150900 | 0.54 |
|  | eat | 757350 | 64 | 2.54 | 00:12:49 | 18:09:41 | 12 | 3824 | 147646 | 0.43 |
|  | sniff | 757350 | 64 | 14.94 | 01:15:26 | 17:07:04 | 0 | 22757 | 128713 | 0.54 |
|  | up | 757350 | 64 | 3.71 | 00:18:44 | 18:03:46 | 6 | 5575 | 145895 | 0.50 |
|  | walk away | 757350 | 64 | 3.89 | 00:19:38 | 18:02:52 | 8 | 5897 | 145573 | 0.50 |

Mouse and rat resident-intruder videos were recorded and annotated at the University of Washington and Washington State University, respectively. CRIM13 is an extensive open-access online catalogue of resident-intruder videos accompanied by detailed annotations of different social and non-social behaviors. The mouse and rat resident-intruder datasets are available to download from the <u>SimBA OSF repository</u>. See the <u>SimBA GitHub repository</u> or the <u>SimBA OSF repository</u> for a detailed depiction of the folder structure and data that is available for download (Fig. 2).

#### Video pre-processing

Video recordings were pre-processed using tools available in SimBA. We shortened, cropped, and saved videos and frames in RGB, greyscale and CLAHE (Contrast-limited adaptive histogram equalization)^48^ enhanced formats at variable frame rates. The CRIM13 dataset was recorded in SEQ file format and converted to MP4 format using SimBA. We also used SimBA to extract frames at specific time-periods, downsample video resolutions, generate gifs and videos. An exhaustive list of SimBA video pre-processing tools and tutorials is available on the <u>SimBA GitHub repository</u>. We used the SimBA to crop, clip, CLAHE enhance, and convert 232 videos in the CRIM13 database^19,47^.

#### Classifier operational definitions

We created a detailed operational definition for each classified behavior. The operational definition described the behavior and its characteristic initiation, duration and end. For the mouse-resident intruder protocol (Table 3), we created operational definitions for 10 predictive classifiers*: attack, pursuit, lateral threat, anogenital sniffing, allogrooming normal, allogrooming vigorous, mounting, scramble, flee, and upright submissive*. For the rat resident-intruder protocol (Table 4), we created operational definitions for 7 predictive classifiers: *attack, anogenital sniffing, lateral threat, approach, boxing, avoidance, and submission*. The CRIM13 dataset was previously annotated for 11 different behaviors: *approach, attack, chase, circle, copulation, drink, eat, sniff, up, clean, and walk away*. For further information, see the <u>CRIM13 website</u>.

**Table 3.**
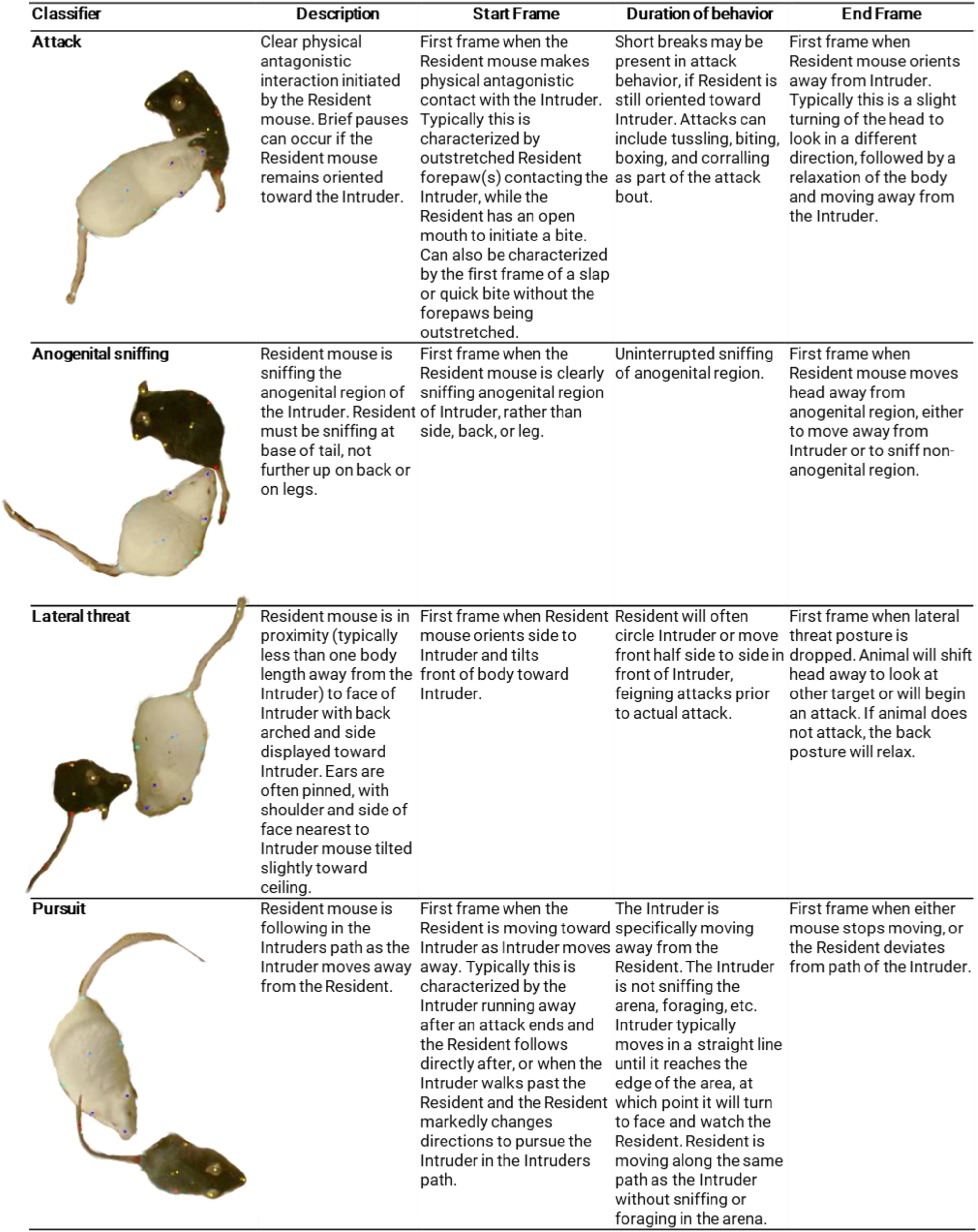
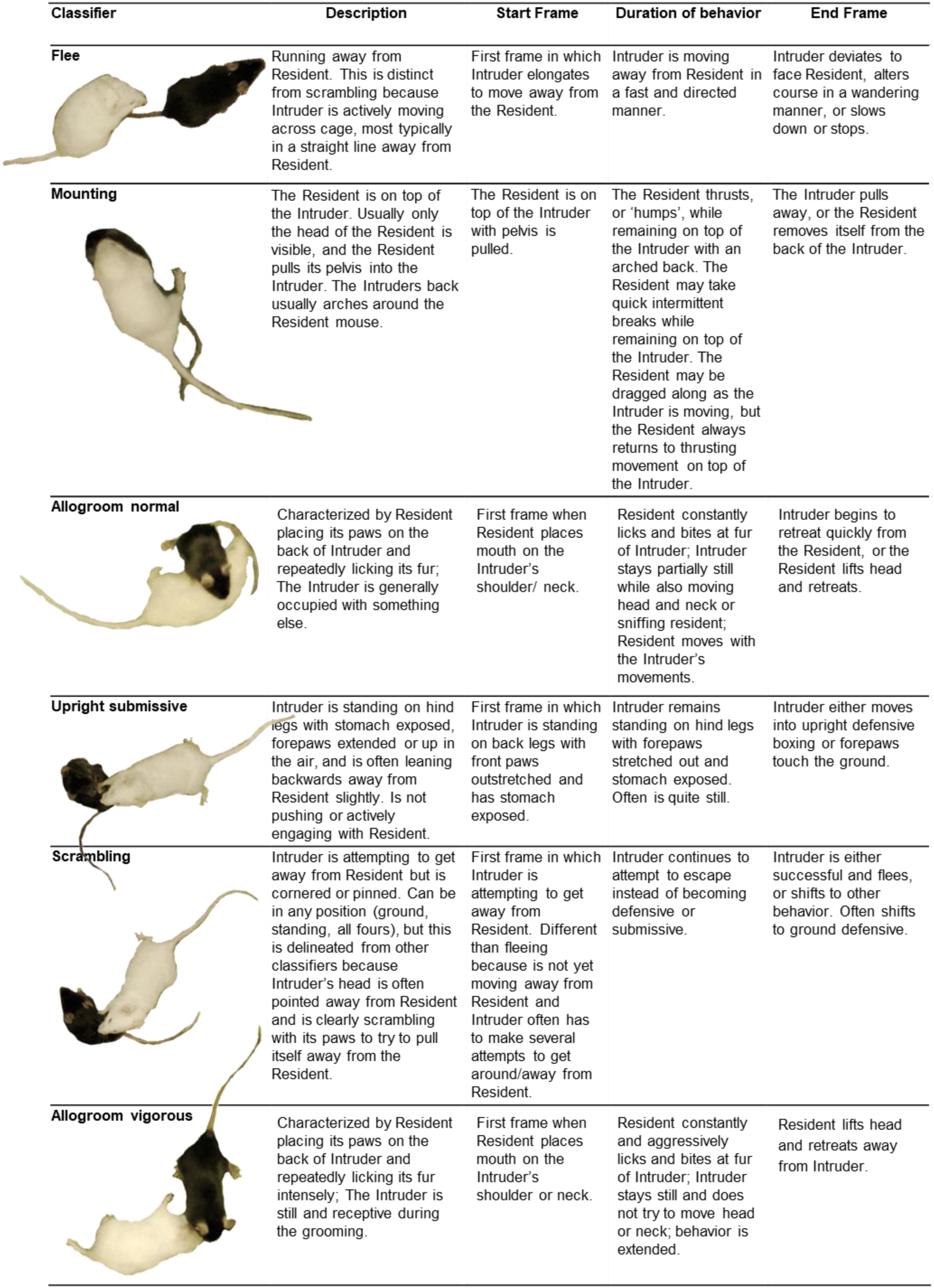
Behavioral operational classifiers: mouse resident-intruder

| Classifier | Description | Start Frame | Duration of behavior | End Frame |
| --- | --- | --- | --- | --- |
| **Attack** | Clear physical antagonistic interaction initiated by the Resident mouse. Brief pauses can occur if the Resident mouse remains oriented toward the Intruder. | First frame when the Resident mouse makes physical antagonistic contact with the Intruder. Typically this is characterized by outstretched Resident forepaw(s) contacting the Intruder, while the Resident has an open mouth to initiate a bite. Can also be characterized by the first frame of a slap or quick bite without the forepaws being outstretched. | Short breaks may be present in attack behavior, if Resident is still oriented toward Intruder. Attacks can include tussling, biting, boxing, and corralling as part of the attack bout. | First frame when Resident mouse orients away from Intruder. Typically this is a slight turning of the head to look in a different direction, followed by a relaxation of the body and moving away from the Intruder. |
| **Anogenital sniffing** | Resident mouse is sniffing the anogenital region of the Intruder. Resident must be sniffing at base of tail, not further up on back or on legs. | First frame when the Resident mouse is clearly sniffing anogenital region of Intruder, rather than side, back, or leg. | Uninterrupted sniffing of anogenital region. | First frame when Resident mouse moves head away from anogenital region, either to move away from Intruder or to sniff non-anogenital region. |
| **Lateral threat** | Resident mouse is in proximity (typically less than one body length away from the Intruder) to face of Intruder with back arched and side displayed toward Intruder. Ears are often pinned, with shoulder and side of face nearest to Intruder mouse tilted slightly toward ceiling. | First frame when Resident mouse orients side to Intruder and tilts front of body toward Intruder. | Resident will often circle Intruder or move front half side to side in front of Intruder, feigning attacks prior to actual attack. | First frame when lateral threat posture is dropped. Animal will shift head away to look at other target or will begin an attack. If animal does not attack, the back posture will relax. |
| **Pursuit** | Resident mouse is following in the Intruder's path as the Intruder moves away from the Resident. | First frame when the Resident is moving toward Intruder as Intruder moves away. Typically this is characterized by the Intruder running away after an attack ends and the Resident follows directly after, or when the Intruder walks past the Resident and the Resident markedly changes directions to pursue the Intruder in the Intruder's path. | The Intruder is specifically moving away from the Resident. The Intruder is not sniffing the arena, foraging, etc. Intruder typically moves in a straight line until it reaches the edge of the area, at which point it will turn to face and watch the Resident. Resident is moving along the same path as the Intruder without sniffing or foraging in the arena. | First frame when either mouse stops moving, or the Resident deviates from path of the Intruder. |

Table 3 continued. Behavioral operational classifiers: mouse resident-intruder
| Classifier | Description | Start Frame | Duration of behavior | End Frame |
| --- | --- | --- | --- | --- |
| **Flee** | Running away from Resident. This is distinct from scrambling because Intruder is actively moving across cage, most typically in a straight line away from Resident. | First frame in which Intruder elongates to move away from the Resident. | Intruder is moving away from Resident in a fast and directed manner. | Intruder deviates to face Resident, alters course in a wandering manner, or slows down or stops. |
| **Mounting** | The Resident is on top of the Intruder. Usually only the head of the Resident is visible, and the Resident pulls its pelvis into the Intruder. The Intruder's back usually arches around the Resident mouse. | The Resident is on top of the Intruder with pelvis is pulled. | The Resident thrusts, or 'humps', while remaining on top of the Intruder with an arched back. The Resident may take quick intermittent breaks while remaining on top of the Intruder. The Resident may be dragged along as the Intruder is moving, but the Resident always returns to thrusting movement on top of the Intruder. | The Intruder pulls away, or the Resident removes itself from the back of the Intruder. |
| **Allogroom normal** | Characterized by Resident placing its paws on the back of Intruder and repeatedly licking its fur; The Intruder is generally occupied with something else. | First frame when Resident places mouth on the Intruder's shoulder/ neck. | Resident constantly licks and bites at fur of Intruder; Intruder stays partially still while also moving head and neck or sniffing resident; Resident moves with the Intruder's movements. | Intruder begins to retreat quickly from the Resident, or the Resident lifts head and retreats. |
| **Upright submissive** | Intruder is standing on hind legs with stomach exposed, forepaws extended or up in the air, and is often leaning backwards away from Resident slightly. Is not pushing or actively engaging with Resident. | First frame in which Intruder is standing on back legs with front paws outstretched and has stomach exposed. | Intruder remains standing on hind legs with forepaws stretched out and stomach exposed. Often is quite still. | Intruder either moves into upright defensive boxing or forepaws touch the ground. |
| **Scrambling** | Intruder is attempting to get away from Resident but is cornered or pinned. Can be in any position (ground, standing, all fours), but this is delineated from other classifiers because Intruder's head is often pointed away from Resident and is clearly scrambling with its paws to try to pull itself away from the Resident. | First frame in which Intruder is attempting to get away from Resident. Different than fleeing because is not yet moving away from Resident and Intruder often has to make several attempts to get around/away from Resident. | Intruder continues to attempt to escape instead of becoming defensive or submissive. | Intruder is either successful and flees, or shifts to other behavior. Often shifts to ground defensive. |
| **Allogroom vigorous** | Characterized by Resident placing its paws on the back of Intruder and repeatedly licking its fur intensely; The Intruder is still and receptive during the grooming. | First frame when Resident places mouth on the Intruder's shoulder or neck. | Resident constantly and aggressively licks and bites at fur of Intruder; Intruder stays still and does not try to move head or neck; behavior is extended. | Resident lifts head and retreats away from Intruder. |

**Table 4.**
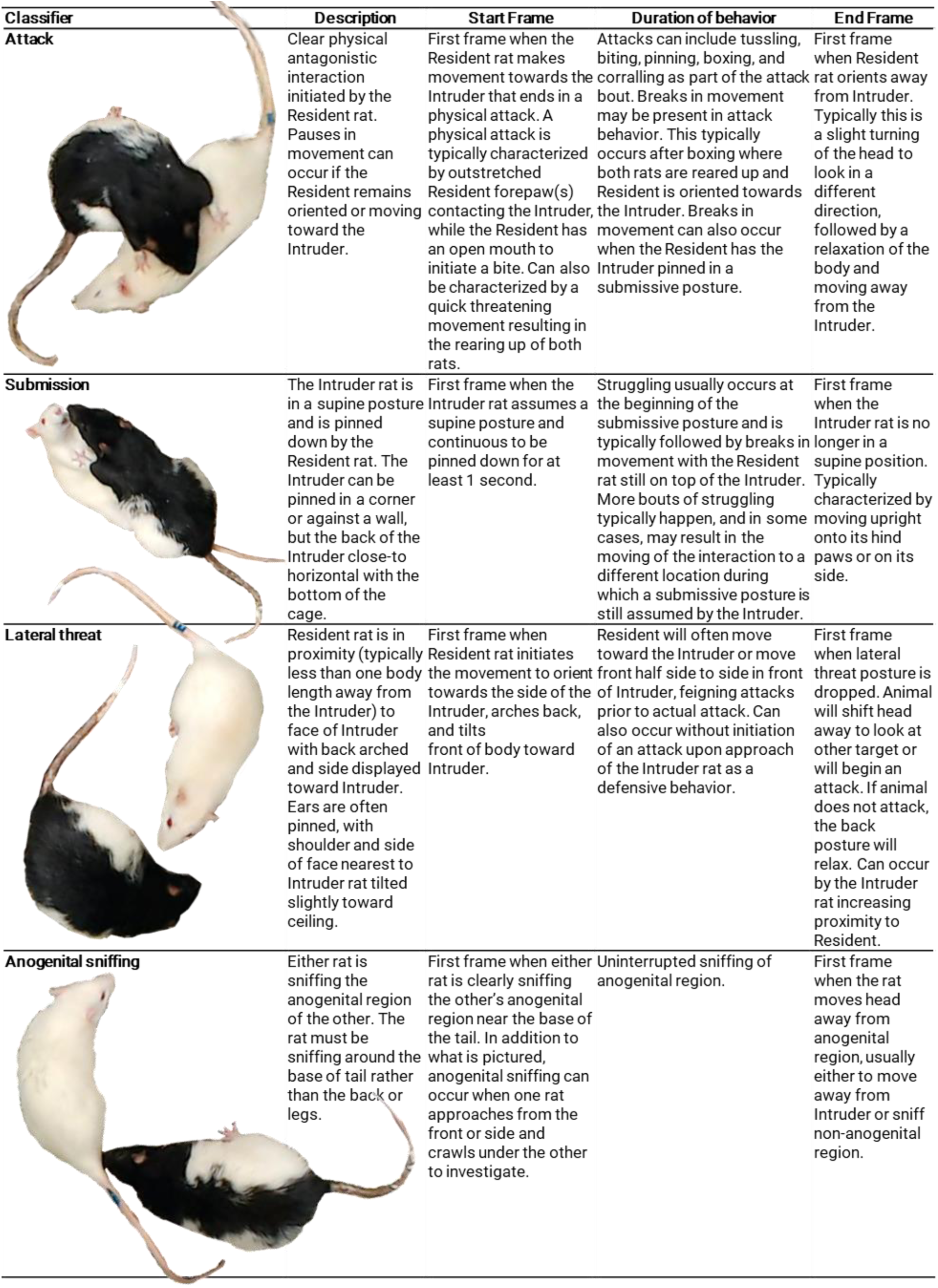
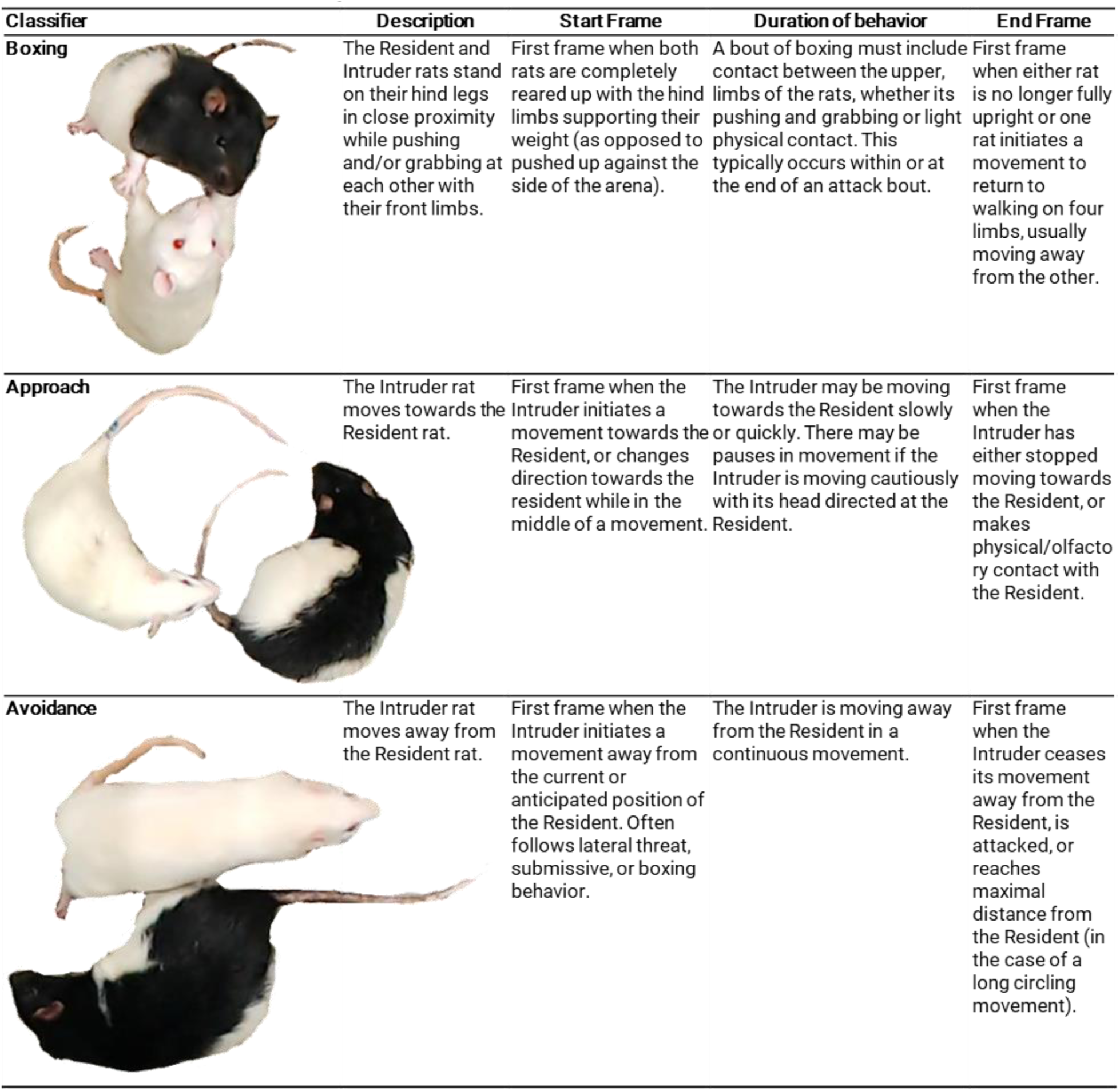
Behavioral operational classifiers: rat resident-intruder

| Classifier | Description | Start Frame | Duration of behavior | End Frame |
| --- | --- | --- | --- | --- |
| **Attack** | Clear physical antagonistic interaction initiated by the Resident rat. Pauses in movement can occur if the Resident remains oriented or moving toward the Intruder. | First frame when the Resident rat makes movement towards the Intruder that ends in a physical attack. A physical attack is typically characterized by outstretched Resident forepaw(s) contacting the Intruder, while the Resident has an open mouth to initiate a bite. Can also be characterized by a quick threatening movement resulting in the rearing up of both rats. | Attacks can include tussling, biting, pinning, boxing, and corralling as part of the attack bout. Breaks in movement may be present in attack behavior. This typically occurs after boxing where both rats are reared up and Resident is oriented towards the Intruder. Breaks in movement can also occur when the Resident has the Intruder pinned in a submissive posture. | First frame when Resident rat orients away from Intruder. Typically this is a slight turning of the head to look in a different direction, followed by a relaxation of the body and moving away from the Intruder. |
| **Submission** | The Intruder rat is in a supine posture and is pinned down by the Resident rat. The Intruder can be pinned in a corner or against a wall, but the back of the Intruder close-to horizontal with the bottom of the cage. | First frame when the Intruder rat assumes a supine posture and continuous to be pinned down for at least 1 second. | Struggling usually occurs at the beginning of the submissive posture and is typically followed by breaks in movement with the Resident rat still on top of the Intruder. More bouts of struggling typically happen, and in some cases, may result in the moving of the interaction to a different location during which a submissive posture is still assumed by the Intruder. | First frame when the Intruder rat is no longer in a supine position. Typically characterized by moving upright onto its hind paws or on its side. |
| **Lateral threat** | Resident rat is in proximity (typically less than one body length away from the Intruder) to face of Intruder with back arched and side displayed toward Intruder. Ears are often pinned, with shoulder and side of face nearest to Intruder rat tilted slightly toward ceiling. | First frame when Resident rat initiates the movement to orient towards the side of the Intruder, arches back, and tilts front of body toward Intruder. | Resident will often move toward the Intruder or move front half side to side in front of Intruder, feigning attacks prior to actual attack. Can also occur without initiation of an attack upon approach of the Intruder rat as a defensive behavior. | First frame when lateral threat posture is dropped. Animal will shift head away to look at other target or will begin an attack. If animal does not attack, the back posture will relax. Can occur by the Intruder rat increasing proximity to Resident. |
| **Anogenital sniffing** | Either rat is sniffing the anogenital region of the other. The rat must be sniffing around the base of tail rather than the back or legs. | First frame when either rat is clearly sniffing the other's anogenital region near the base of the tail. In addition to what is pictured, anogenital sniffing can occur when one rat approaches from the front or side and crawls under the other to investigate. | Uninterrupted sniffing of anogenital region. | First frame when the rat moves head away from anogenital region, usually either to move away from Intruder or sniff non-anogenital region. |

**Table 4 continued. Behavioral operational classifiers: rat resident-intruder**
| Classifier | Description | Start Frame | Duration of behavior | End Frame |
| --- | --- | --- | --- | --- |
| **Boxing** | The Resident and Intruder rats stand on their hind legs in close proximity while pushing and/or grabbing at each other with their front limbs. | First frame when both rats are completely reared up with the hind limbs supporting their weight (as opposed to pushed up against the side of the arena). | A bout of boxing must include contact between the upper, limbs of the rats, whether its pushing and grabbing or light physical contact. This typically occurs within or at the end of an attack bout. | First frame when either rat is no longer fully upright or one rat initiates a movement to return to walking on four limbs, usually moving away from the other. |
| **Approach** | The Intruder rat moves towards the Resident rat. | First frame when the Intruder initiates a movement towards the Resident, or changes direction towards the resident while in the middle of a movement. | The Intruder may be moving towards the Resident slowly or quickly. There may be pauses in movement if the Intruder is moving cautiously with its head directed at the Resident. | First frame when the Intruder has either stopped moving towards the Resident, or makes physical/olfactory contact with the Resident. |
| **Avoidance** | The Intruder rat moves away from the Resident rat. | First frame when the Intruder initiates a movement away from the current or anticipated position of the Resident. Often follows lateral threat, submissive, or boxing behavior. | The Intruder is moving away from the Resident in a continuous movement. | First frame when the Intruder ceases its movement away from the Resident, is attacked, or reaches maximal distance from the Resident (in the case of a long circling movement). |

#### Resident-intruder protocol

For the mouse resident-intruder protocol, we recorded dyadic encounters between two male mice (cage size: 28×17×14cm) or two female mice (cage size: 28×19×12cm) in clear polycarbonate home-cages with fresh bedding, or two male mice in Med-Associates operant chambers (chamber size: 21×18×13cm) using protocols detailed elsewhere^49,50^. The intruder was a black coat-coloured C57BL/6J mouse (Jackson Labs, #000664), and the resident was a white coat-coloured CD-1 mouse (Charles River, #022). For the rat resident-intruder protocol, we recorded resident-intruder dyadic encounters between a male intruder Sprague Dawley rat (Simonsen Laboratories) and male resident Long-Evans rat (Simonsen Laboratories) in clear polycarbonate cages with fresh bedding (cage size: 33×46×19cm) using a previously detailed protocol^51^.

#### Video recordings

Male mice in were recorded at 30-80fps with USB3.0 cameras (acA2040-120uc - Basler ace, Basler) using fixed-focal length lenses (Edmund Optics, NJ, 16mm/F1.4) from a 90° angle at variable resolutions (W:1000-1200px, H:1255-2056px) using the pylon camera software (Basler). Male rats were recorded at 60fps and 1280×720 resolution at a 90° angle from above using a Logitech C922 USB web-camera. Importantly, all recorded videos used to build behavioral classifiers were re-sampled in SimBA to 30fps before being used to generate machine learning classifiers. All video recordings can be found on the <u>SimBA OSF repository</u>. The Caltech Resident-Intruder Mouse (CRIM13) dataset was recorded at 25fps and 640×480 resolution from a 90° angle.

#### Project management

SimBA has separate drop-down toolbar menus for creating and loading animal tracking projects and predictive behavior classification projects. Animal tracking data is generated within animal tracking projects and subsequently exported to predictive classification projects. The tracking data is used to create predictive classifiers or analyzed using previously generated predictive classifiers. Within the classification project menus, the user can also analyze animal movements and behavioral patterns based on <u>user-drawn region-of-interests (ROIs)</u>.

#### Animal pose-estimation tracking

DeepLabCut and DeepPoseKit projects were accessed and managed through the Tracking▷DeepLabCut and Tracking▷DeepPoseKit toolbars of SimBA. We used the DeepLabCut and DeepPoseKit labeling interfaces to annotate eight body-parts on each of the two animals (Fig. 3a). For alternative protocols, SimBA supports any body labeling schematic if first defined by the user in the Create Project menu of SimBA (Fig. 3c-d). All generated tracking models, together with the annotated images, are available for download on the <u>SimBA OSF repository</u>. The <u>SimBA OSF repository</u> also contains annotations and models for detecting black and white coat-colored mice through alternative convolutional learning architectures (e.g., mask RCNN^52^, YOLO^53^).

**Figure 3.**
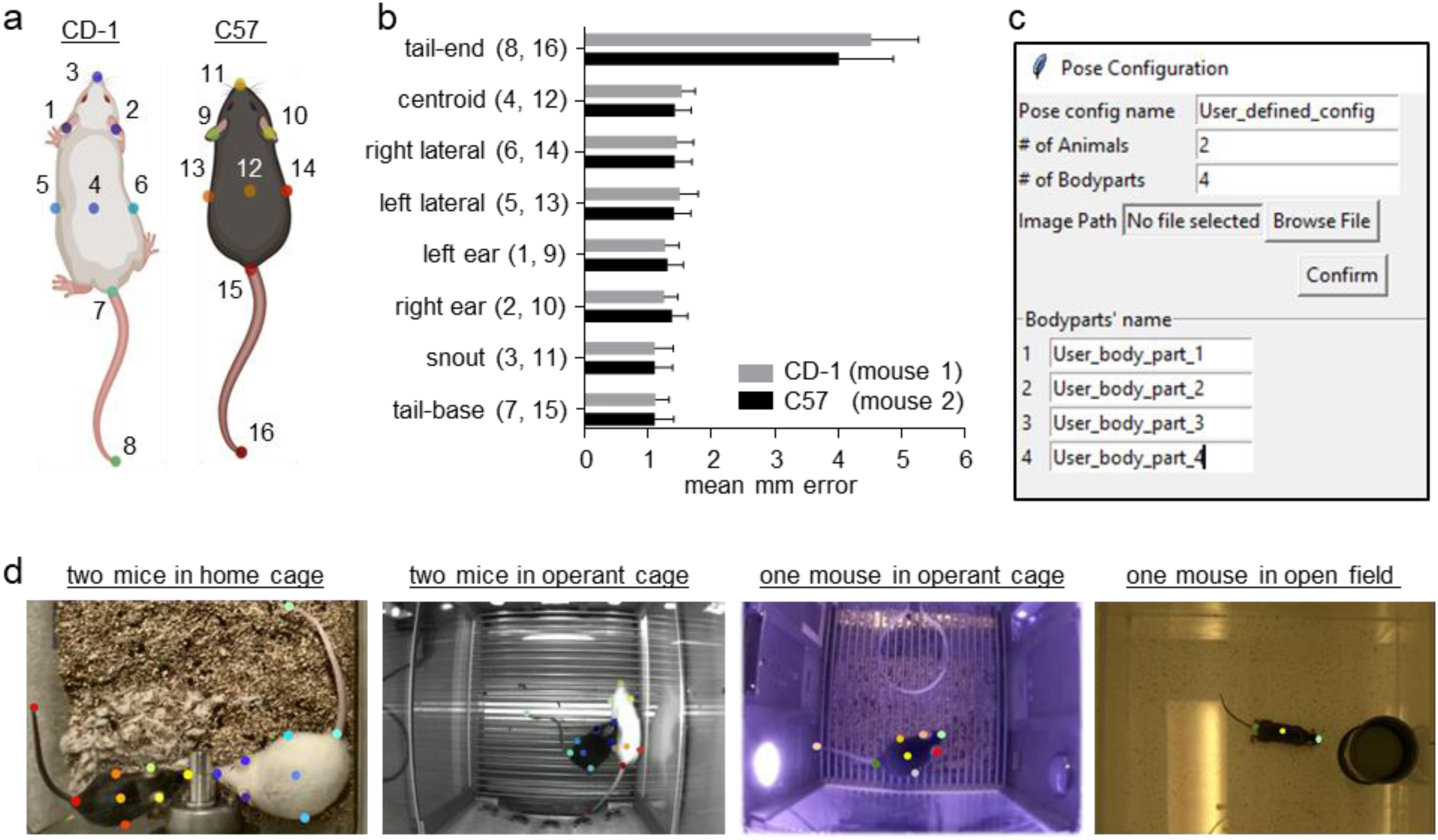
Animal pose-estimation body-part tracking. (a) Schematic representation of the 16 body-parts annotated and tracked in the resident-intruder protocols using DeepLabCut and DeepPoseKit. The adjacent numbers represent the order which the body-parts were annotated in the DeepLabCut and DeepPoseKit annotation interfaces. (b) The mean millimeter tracking errors for the body-parts in the mouse RGB residentintruder model. (c) The SimBA *flexible annotation module* for creating and importing user-defined body-part tracking schemas for generation of supervised machine learning classifiers. (d) Examples of body-part tracking schemas created in SimBA for pose-estimation and supervised machine learning classifiers.

### Behavior classifications

#### Social classifiers

We created and managed machine classification projects using the File▷ create project menu and File▷Load project menu in SimBA. We imported the pose-estimation tracking data in CSV file format together with the accompanying videos into each project. We used SimBA to extract the individual frames for each of the videos in the projects (*note*: extraction of frames is a CPU-intensive and time-consuming process that depend on the frame-rate, length, and resolution of the videos). Creating frames for individual videos, however, is only necessary for annotating behaviors and creating new classifiers in SimBA and is not necessary for visualizing and analyzing videos. See the <u>SimBA GitHub repository</u> for tutorials and recommended approaches for extracting frames in different use-case scenarios.

#### Time and distance standardization

We used SimBA to standardize distance (pixel/mm) and time (frames per second) measures across different video recordings. SimBA allows the user to define and draw a known distance in the video (for example, the arena length or the arena width) and this distance is used to convert Euclidean pixel distances to millimeter distances. Furthermore, SimBA automatically registers the frame rate of each video, and this information is was used to standardize calculations across rolling time windows.

#### Correcting inaccurate tracking

We designed two outlier correction tools that identify and correct gross poseestimation tracking inaccuracies by detecting outliers based on movements and locations of body-parts in relation to the animal body-length (Fig. 4a). To compute an initial reference value, SimBA first calculates the mean Euclidian millimeter distance between two user-defined body-parts across the entire recording (*L*). We set the body-parts to be the nose and the tail-base. We also set two criterion values, one criterion value for ‘movement outliers’ (*V_1_*) and one criterion value for ‘location outliers’ (V_2_). A body-part coordinate was first corrected as a ‘movement outlier’ if the movement in Euclidean pixels of the body-part, across two sequential frames, was equal or more than *L* × V_1_ (Fig. 4b, top). If a body-part was found to be a movement outlier in six consecutive frames a new ground truth value was sampled. Second, a body-part coordinate was corrected as a ‘location outlier’ if the distance in Euclidian millimeters of that body-part, to at least two other body-parts belonging to the same animal (excluding the tail-end), was equal or more than *L* × V2 (Fig. 4b, bottom). Body-parts detected as outliers were corrected to their last reliable coordinate. The SimBA outlier correction tools generate logs of the total number of corrected body-parts, and ratio of corrected body-parts, in each video of the project.

**Figure 4.**
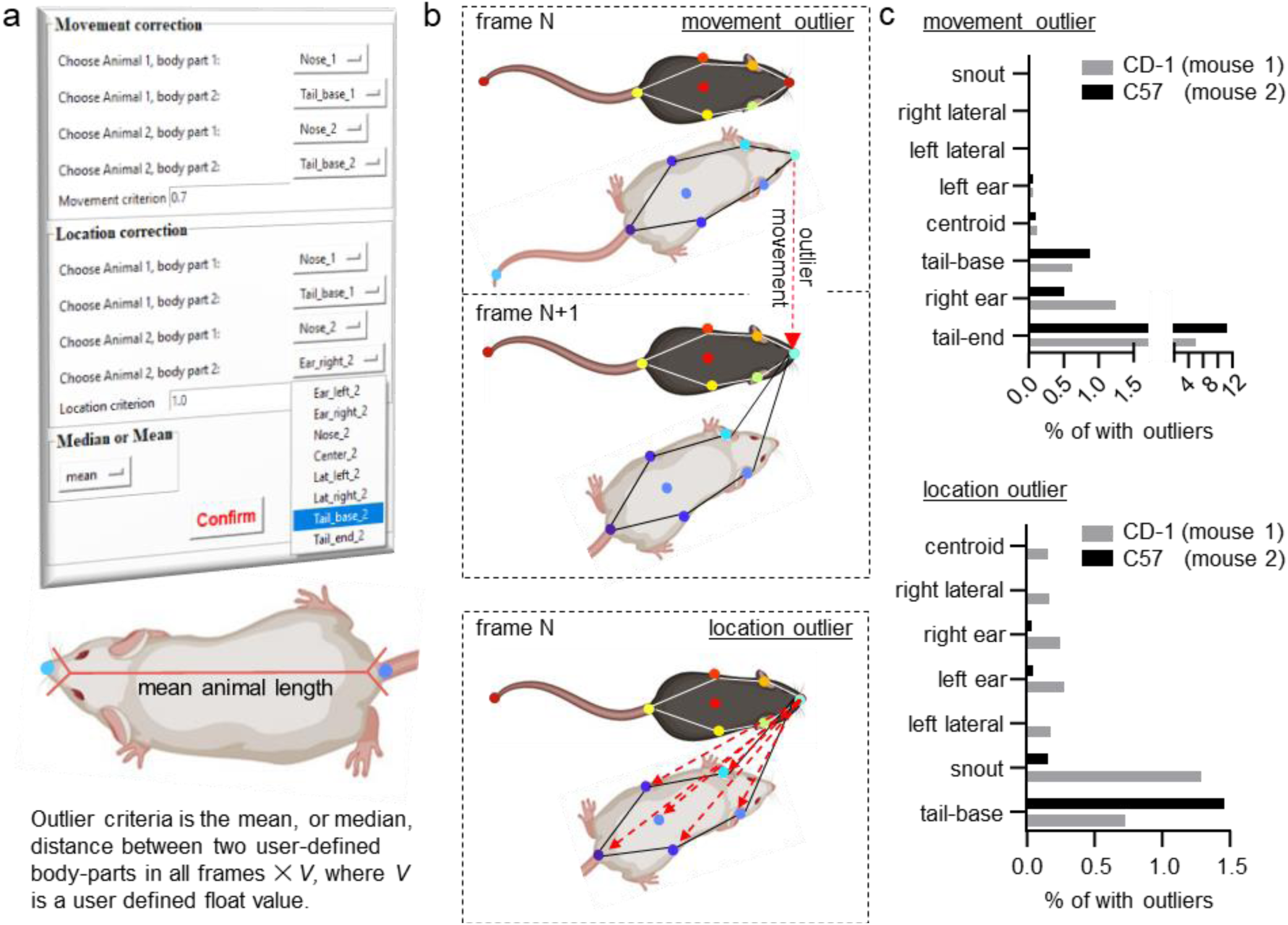
Tools in SimBA for removing inaccuracies of pose-estimation body-part tracking. (a) SimBA calculates the mean or median distance between two user-defined body-parts across the frames of each video. We set the user-defined body-parts to be the nose and the tail-base of each animal. The user also defines a *movement criterion value*, and a *location criterion value*. We set the movement criterion to 0.7, and location criterion to 1.5. Two different outlier criteria are then calculated by SimBA. These criteria are the mean length between the two user-defined body parts in all frames of the video, multiplied by the either user-defined *movement criterion value* or *location criterion value*. SimBA corrects movement outliers prior to correcting location outliers. **(b)** Schematic representations of a pose-estimation body-part ‘movement outlier’ (top) and a ‘location outlier’ (bottom). A body-part violates the movement criterion when the movement of the body-part across sequential frames is greater than the movement outlier criterion. A body-part violates the location criteria when its distance to more than one other body-part in the animals’ hull (except the tail-end) is greater than the location outlier criterion. Any body part that violates either the movement or location criterion is corrected by placing the body-part at its last reliable coordinate. **(c)** The ratio of body-part movements (top) and body-part locations (bottom) detected as outliers and corrected by SimBA in the RGB-format mouse resident-intruder data-set. For the outlier corrected in rat and the CRIM13 datasets, see the <u>SimBA GitHub repository</u>.

#### Decision tree features

SimBA combines the registered distance (pixels/mm) and time (frames per second) variables for each video with the corrected tracking data to calculate an exhaustive list of distance, movement, angles, areas, and paths metrics and their deviations and rank for individual frames and across rolling windows^24^. The features calculated by SimBA, and how many features that are calculated, depend on the body-parts tracked during pose-estimation. When using a 16-body-part tracking schematic on two animals, SimBA extracts a set of 498 different metrics relevant for classifying social and non-social behaviors in the mouse and rat resident datasets. When using other pre-defined body-part configurations, or user-defined body-part configuration, SimBA calculates a reduced and/or generic feature set based on all individual body-parts and their relationships and movements. Lists and descriptions of the different features calculated in different use-case scenarios are available on the <u>SimBA GitHub repository</u>.

#### Behavior annotations

SimBA has an in-built event logger that displays individual frames alongside the video. Within the event logger interface, the user selects the behavioral events (defined during project creation) that are occurring in the displayed frames, and SimBA concatenates the logged events with the features calculated from the corrected pose-estimation tracking data. We used the SimBA event logger to annotate frames as containing, or not containing, the behaviors of interest in the mouse and rat resident-intruder videos. A tutorial for using the SimBA event logger is available on the <u>SimBA GitHub repository</u>. In many alternative scenarios, the behavioral events may have already been logged through alternative software tools (e.g., JWatcher, Noldus Observer). In these scenarios it is preferable to build predictive classifiers from previously generated event logs without performing further annotations in SimBA. The exact code will depend on the structure of the previously generated event logs. For example, the CRIM13 dataset was annotated using a MATLAB annotation tool^54^. We concatenated the CRIM13 annotations to the CRIM13 tracking data using a <u>modifiable script</u> that can be downloaded from the <u>SimBA GitHub repository</u>. We request that individuals with already-annotated datasets contact us for developing scripts to easily import annotations into SimBA.

#### Classification ensembles

Random forest classifiers are intuitive algorithms generated by majority verdicts from decision trees that split the data along feature values to separate distinct classes (e.g., *attack* or *not attack)^55^*. We used SimBA to create random forest classifiers, with one classifier for each of the behaviors of interest. Each classifier contained 2k decision trees. The number of trees was chosen as a large, computationally feasible, amount^56^. Random forest classifiers accept a range of *hyperparameters* that specify how the decision trees should be generated. SimBA accepts a range of different random forest hyperparameter settings, but users unfamiliar with the available parameters can import recommended or previously successful settings based on classifiers of behaviors with similar frequency and salience. Hyperparameter meta files, that can be imported into SimBA and contain the hyperparameters used to generate the current classifiers, can be downloaded from the <u>SimBA OSF repository</u>. We evaluated each classifier by calculating its *precision, recall*, and *F1-scores* after 5-fold shuffle cross-validations on 20% of the datasets annotated in the SimBA event logger. Precision was calculated as

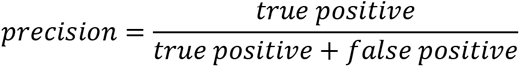

and denotes the proportion of frames correctly classified as displaying the behavior of all the frames classified as containing the behavior. Recall was calculated as

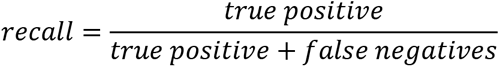

and denotes the proportion of all the frames containing the behavior that were correctly classified as containing the behavior. F1 was calculated as

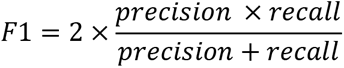

and is the harmonic mean of the precision and recall scores.

Random forest classifiers - as other classification techniques - are sensitive to class imbalances. Class imbalances are prevalent in most datasets and occur when observations of the majority class (i.e., observations of the *absence* of the classified behavior) substantially outweigh observations of the minority class (i.e., observations of the *presence* of the classified behavior). For example, in the current mouse resident intruder dataset *flee* events were present in 1% of the annotated frames (Table 2). In this scenario the random forest decision trees can classify most *flee* observations correctly by disregarding the presence of *flee* events. To prevent this, SimBA supports several class re-balancing methods, including random under-sampling of the majority class, and over-sampling of the majority class using SMOTE^57^ or SMOTE-EEN^58^. We maximized the F1-score of the classifiers by random under-sampling the majority class (i.e., observations of the absence of the classified behavior) at ratios up to 1:46. See Table 2 for the random under-sampling ratios used for each classifier.

#### Classifier evaluations

We used SimBA to generate classifier learning curves. In these learning curves we evaluated F1-scores after performing 5-fold cross-validations using 1, 25, 50, 75 and 100% of the shuffled data sets to predict the classified behaviors on 20% of the datasets. Learning curves indicate how inclusion of further logged behavioral events affect classifier performance. Furthermore, we used SimBA to generate precision-recall curves^59^, that visualize how classifiers can be titrated to balance the sensitivity versus the specificity of the classifications^60^ through different discrimination thresholds. We also used SimBA to calculate *mean decrease gini impurity* and *permutation importance* scores which gauge the importance of individual features for correctly classifying individual behaviors^55^. *Mean decrease gini impurity score* or *impurity importance* represents the average gain in purity of a decision tree when the feature is included. *Feature permutation importance* represents the loss of predictive power when the feature, and no other feature, is scrambled. Lastly, we evaluated the F1 scores of each classifier through 5-fold cross-validations after shuffling the behavioral event annotations made by the experimenter in the training set, and testing the classifier on the un-shuffled, correctly annotated behavioral data^61^. The final random forest classifiers for mouse and rat resident-intruder protocols, and the CRIM13 dataset, can be downloaded through the <u>SimBA OSF repository</u> accessible through the <u>SimBA GitHub repository</u>.

#### Procedural runtimes

We used the python timeit module to measure the execution time of six important SimBA scripts on two computers with different hardware specifications (Fig. 5). We also measured the mean time to extract 1k individual image frames from a video recorded at six different resolutions on the two computers. We executed each script was five times and recorded the mean execution time.

**Figure 5.**
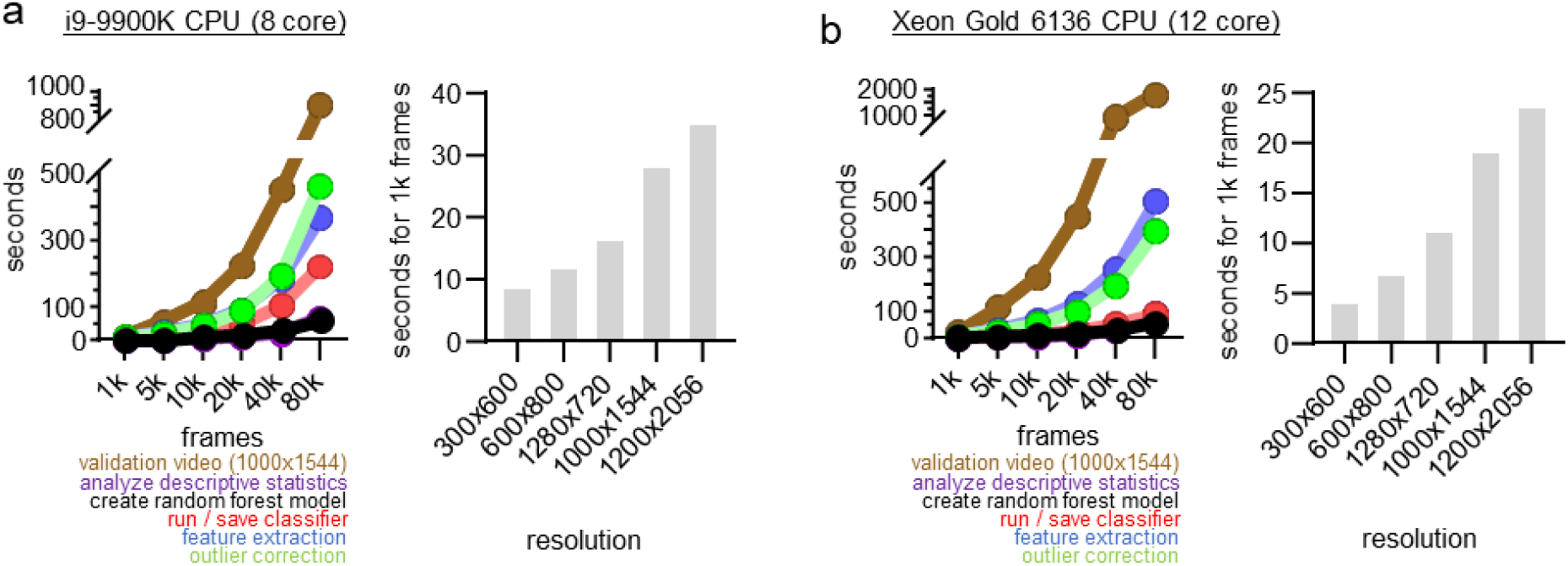
Approximate procedural runtimes for processing different sized data-sets in SimBA. (a) an 8-core Intel i9 CPU and (b) a 12-core Xeon Gold CPU. Time in seconds to perform outlier corrections, feature extraction using 16 body-parts, generating a random forest with 2k trees, performing / saving machine learning classifications, calculating descriptive statistics of machine classifications, generating a validation video, and extracting individual image frames from a video recorded at six different resolutions (videos recorded at 975kbps bitrate). *Note*: only runtimes for creating a validation video and extracting frames depend on the resolution of the videos. See the <u>SimBA GitHub repository</u> for more information.

## RESULTS

### Animal tracking

We used the DeepLabCut^27^ interface to label 16 body-parts on a male CD-1 and C57BL/6J mouse in 6765 frames from 50 videos in the resident-intruder protocol recorded in RGB format (Fig. 3a). We labeled a further 1550 frames of female mice in the resident-intruder protocol (kindly shared by Emily Newman, Tufts). The total number of frames annotated was 8315. The mean tracking error in millimeters for each of the 16 body-parts in the mouse home-cage resident intruder protocol, recorded in RGB format, is shown in Fig. 3b. For most body-parts the mean Euclidean tracking error was around 1 mm. The tail-ends were associated with larger mean tracking errors between 4-5 mm.

We used the network weights generated from the 8315 annotated images to create additional body-part tracking models for alternative experimental protocols. For these additional tracking models, we labelled a further 800 frames from home-cage environments with male rats, and 3200 CLAHE enhanced images from the Caltech Resident-Intruder Mouse (CRIM13) dataset^19,47^. All annotations were converted to greyscale and CLAHE enhanced formats, and models were generated based on these annotations. All annotated images, and tracking models, can be downloaded from the <u>SimBA OSF repository</u>.

### Procedural Runtime Performance of SimBA functions

The computation speed of several important SimBA functions are shown in Fig. 5. For 1-80k frames on the 8-core Intel i9 CPU-supported computer, pose-estimation outlier correction was performed in 10-465s, feature extraction 5-367s, random forest generation 0-68s, running/saving classifications 4-222s, calculating descriptive statistics 0-67s, and creating a validation video took 11-906s (Fig. 5a). For 1-80k frames on the 12-core Intel Xeon Gold CPU-supported computer, pose-estimation outlier correction was performed in 4-394s, feature extraction 6-504s, random forest generation 0-53s, running/saving classifications 3-90s, calculating descriptive statistics 0-89s, and creating a validation video took 22-1804s (Fig. 5b). Current SimBA efforts are focused on accessible creation of social behavior machine learning classifiers, and future efforts will optimize execution runtimes.

### Social Classifiers

#### Outlier corrections

We used SimBA to correct gross pose-estimation tracking inaccuracies based on impossible locations and movements of animal body-parts (Fig 4a-b). In accordance with the tracking data, the tail-ends represented most of the corrected body-parts based on body-part movements (Fig. 4c). The tail-bases and snouts represented most of the corrected body-parts based on body-part locations. In total, 0.013% of all body-parts were corrected following the movements outlier correction, and a further 0.006% of the body parts were corrected following the location correction. For tracking inaccuracies corrected in other data-sets, see the <u>SimBA GitHub repository</u>.

#### Classifier datasets

We used the SimBA event logger to annotate video frames as containing, or not containing, the behavior. The number of annotated video frames and the percent of video frames that were annotated as containing the behavior are shown in Table 2. For the classifiers in the mouse resident-intruder dataset, we annotated between 103k and 472k frames. The classified behaviors were annotated as present in 0.73-6.57 % of the video frames. For the predictive classifiers in the rat resident-intruder dataset, we annotated 136k frames and the classified behaviors were annotated as present in 2.48-16.40 % of the frames. Of the original 232 videos in the CRIM13 dataset, we identified 67 videos and 842k frames that contained social-interactions between nonanesthetized black and white coat-colored animals without the presence of a human. We used 64 of the videos and 757k in the training and test sets and saved three videos for validation. The classified behaviors had been annotated as present in 0.36-14.94 % of the video frames.

#### Mouse resident-intruder classifier performances

Mean precision, recall and F1-scores for the presence and absence of the behaviors following 5-fold shuffle cross-validation of the final classifiers in the mouse resident-intruder protocol is shown in Fig. 6. Frame-wise classification performance for the *presence* of the behaviors as measured by F1 were 0.778-0.983, precision between 0.899-0.997, and recall between 0.685-0.972. Frame-wise classification performance scores for the *absence* of the behaviors were 0.982 or higher (Fig. 6c). Five-fold cross validation learning curves using 1-100% of the annotated data (Fig. 6a) showed that the number of annotated images positively correlate with F1 score. Precision-recall curves (Fig. 6b) indicated optimal classifier performance as measured by F1 at discrimination thresholds between 0.41-0.79.

**Figure 6.**
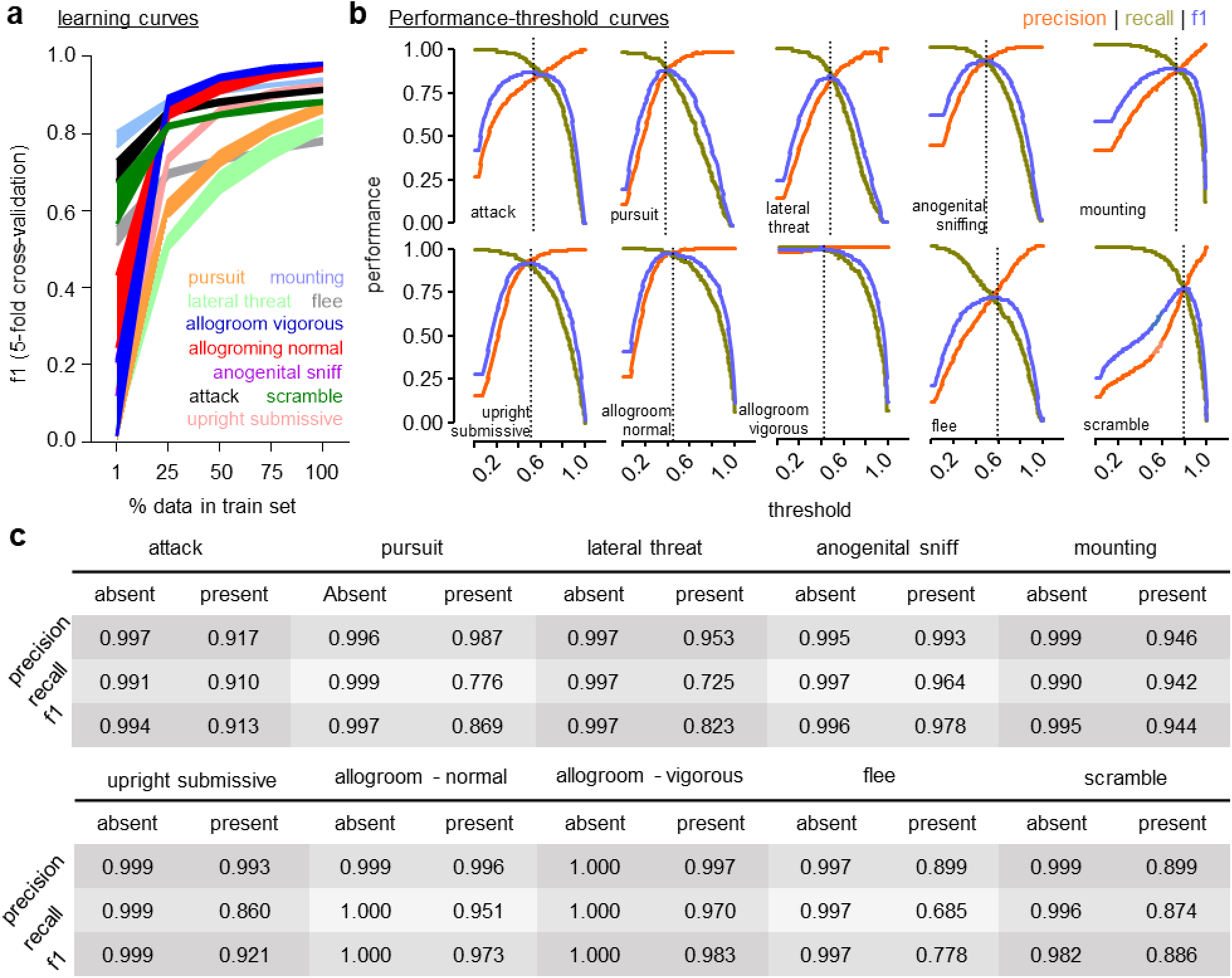
Evaluations of mouse resident-intruder predictive classifiers. (a) Learning curves were created using 2k trees, 5 data splits (1-100%), and with shuffled 5-fold cross-validation at each data split. Errors represent ± SEM. (b) Classification precision, recall, and F1 scores at different discrimination thresholds. The dotted line represents the discrimination threshold at maximal F1 score. See Table 1 for the optimal thresholds for each classifier. (c) Mean classifier precision, recall, and F1 score evaluated by shuffled 5-fold shuffle cross-validation. See Methods for equations and detailed descriptions of precision, recall, and f1 metrics.

#### Rat resident-intruder classifier performances

Mean precision, recall and F1-scores for the presence and absence of the behaviors following 5-fold shuffle cross-validation of the final classifiers in the mouse residentintruder protocol is shown in Fig. 7. Frame-wise classification performance for the *presence* of the behaviors as measured by f1 were 0.918-0.985, precision between 0.990-0.997, and recall between 0.855-0.987. Frame-wise classification performance scores for the *absence* of the behaviors were 0.994 or higher (Fig. 7c). Five-fold cross validation learning curves using 1-100% of the annotated data (Fig. 7a) showed that the number of annotated images positively correlate with f1-score. Precision-recall curves (Fig. 7b) indicated optimal classifier performance as measured by f1 at discrimination thresholds between 0.45-0.55.

**Figure 7.**
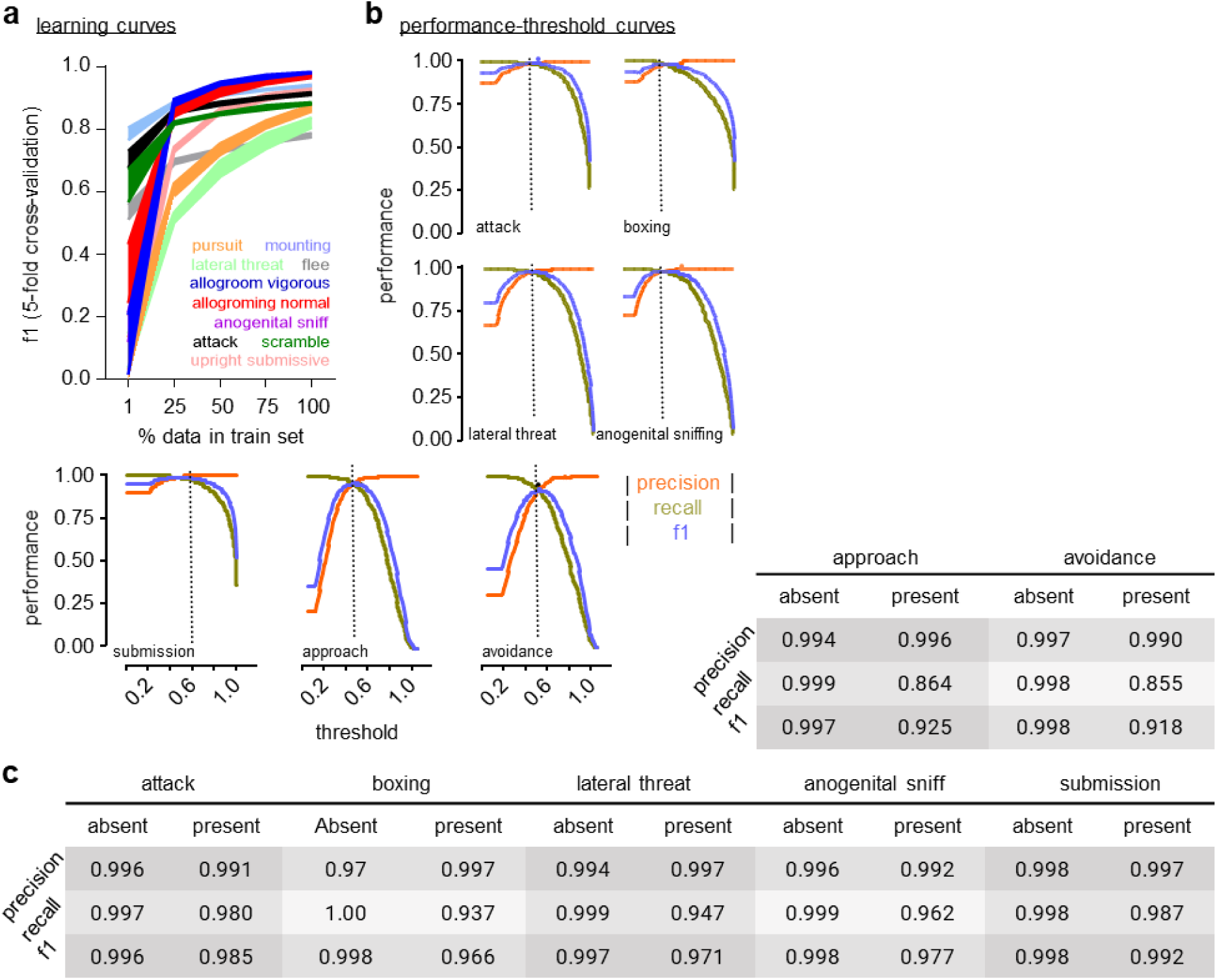
Evaluations of rat residentintruder predictive classifiers. (a) Learning curves were created using 2k trees, 5 data splits (1-100%) and with shuffled 5-fold cross-validation at each data split. Errors represent ± SEM. (b) Classification precision, recall, and F1 scores at different discrimination thresholds. The dotted line represents the discrimination threshold at maximal F1 score. See Table 3 for the optimal threshold for each classifier. (c) Mean classifier precision, recall, and F1 score evaluated by shuffled 5-fold shuffle cross-validation. See Method for equations and descriptions of precision, recall, and F1 metrics.

#### CRIM resident-intruder classifier performances

Mean precision, recall and F1-scores for the presence and absence of the behaviors following 5-fold cross-validation of the final classifiers in the mouse resident-intruder protocol is shown in Fig. 8. Frame-wise classification performance for the *presence* of the behaviors as measured by F1 were 0.739-0.957, precision between 0.910-0.997, and recall between 0.639-0.919. Frame-wise classification performance scores for the *absence* of the behaviors were 0.991 or higher (Fig. 8c). Five-fold cross validation learning curves using 1-100% of the annotated data (Fig. 8a) showed that the number of annotated images positively correlate with F1-score. Precision-recall curves (Fig. 8b) indicated optimal classifier performance as measured by F1 at discrimination thresholds between 0.40-0.58.

**Figure 8.**
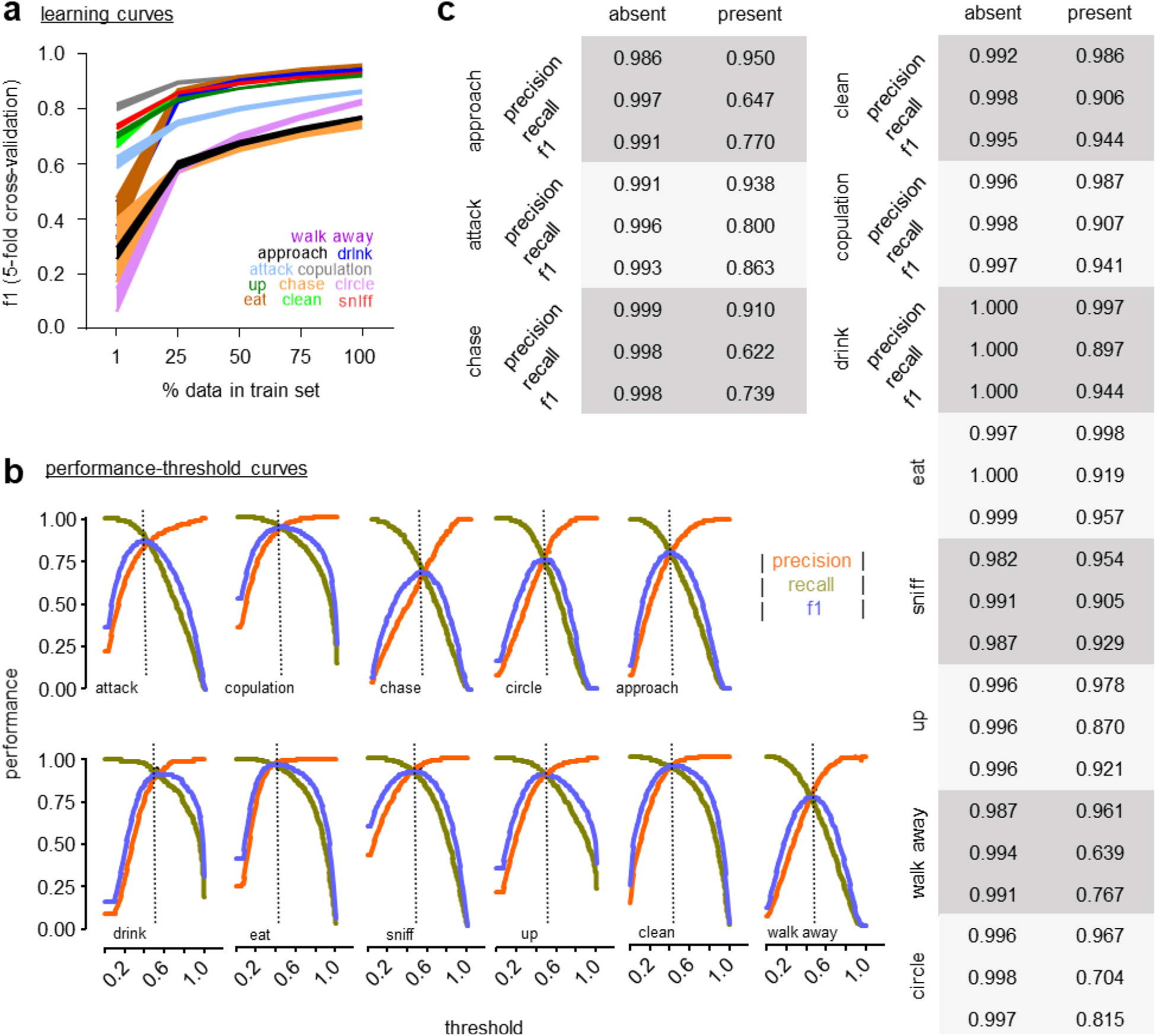
Evaluations of CRIM13 mouse resident-intruder predictive classifiers. (a) Learning curves were created using 2k trees, 5 data splits (1-100%), and with shuffled 5-fold cross-validation at each data split. Errors represent ± SEM. (b) Classification precision, recall, and F1 scores at different discrimination thresholds. The dotted line represents the discrimination threshold at maximal F1 score. See Table 3 for the optimal threshold for each classifier. (c) Mean classifier precision, recall, and F1 evaluated by shuffled 5-fold shuffle cross-validation. See Method for equations and descriptions of precision, recall, and F1 metrics.

#### Feature permutation importances

The four most significant features for each classified behavior, as measured by permutation importance’s, are shown in Fig. 9. Feature permutation importances represents the performance loss when the specific feature is scrambled. Greater permutation importance values represent a higher feature significance for correct classifications. For complete lists of feature significances for each classified behavior as measured by *mean decrease gini impurity* and permutation importance’s, see the <u>SimBA OSF repository</u>. For example, *attacks* in the mouse resident-intruder dataset (Fig. 9a) were classified - in part - based on the amount of movements, the percentile rank of movements / deviation of movements relative to the mean movements in the video. Attack classifications in the rat resident-intruder dataset were also, in part, based on amount of movements, but also the presence of reduced pose-estimation detection probabilities and decreased metric distances between the body-parts in the animal hulls. This can be explained by rats, but not mice, typically assuming stable close-contact *rearing* postures during attack bouts that decrease pixel distances between bodyparts and may cause occlusion of individual body-parts.

**Figure 9.**
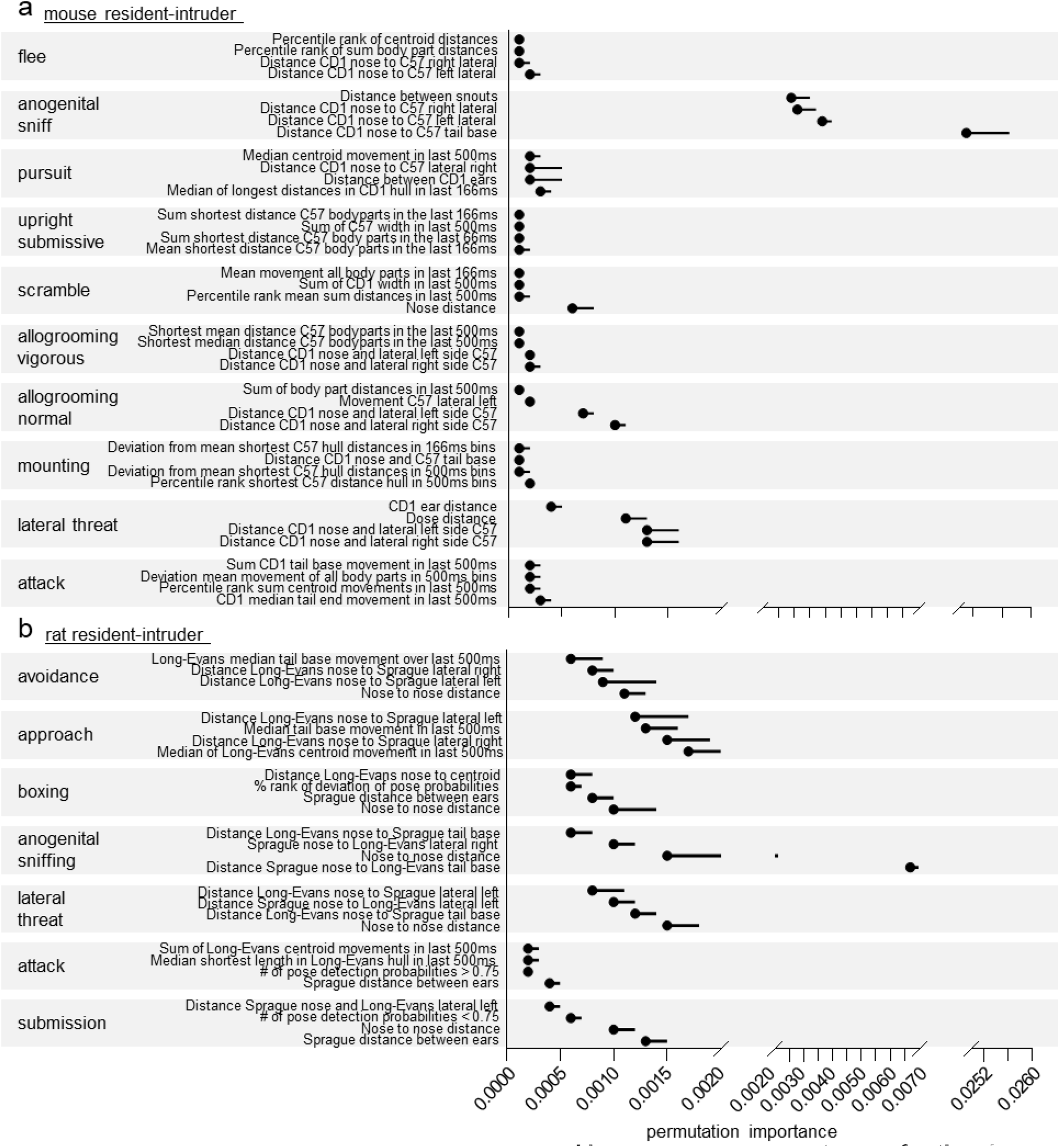
The four most significant features as measured by permutation importances for the classifiers. (a) mouse resident intruder and (b) rat resident-intruder data-sets. Feature permutation importance represents the performance classification degradation when the specific feature, and no other feature, is scrambled. A complete list of feature permutation and gini importance’s are available through the <u>SimBA OSF repository</u>.

#### Annotation scrambling

We performed 5-fold cross-validations where we randomly shuffled the human-made behavior annotations in the training set, before testing the classifier generated from the shuffled data on the correctly annotated, non-shuffled, training set^61^. Shuffling the annotations in the training sets to 0.00-0.0036 decreased F1-scores (Fig. 10).

**Figure 10.**
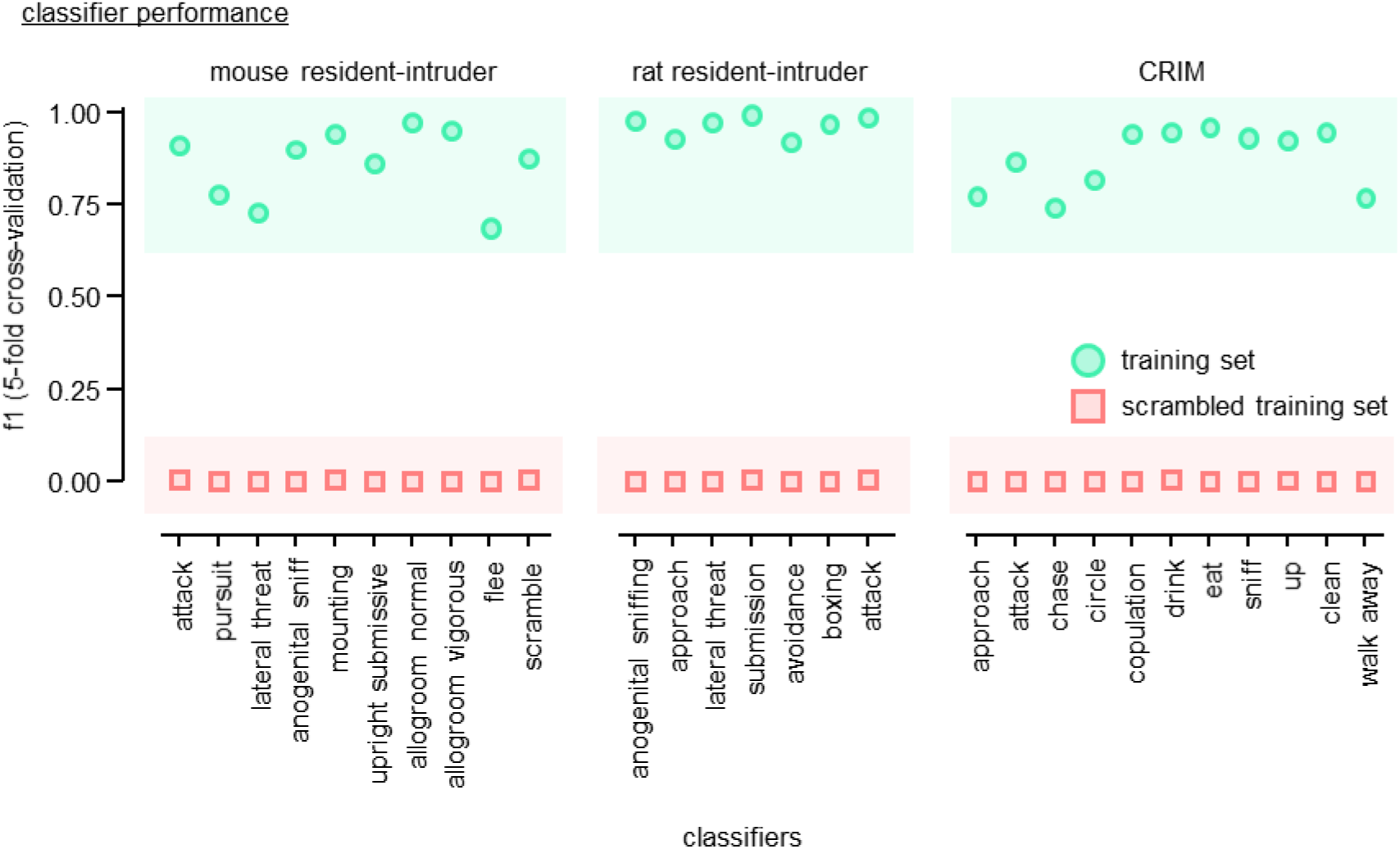
Classifier performance after randomly scrambling the human-made annotations in the training set. Performance was evaluated as F1-score for the presence of the target behavior, measured by shuffled 5-fold cross-fold validation after randomly scrambling the human annotations in the training set. The classifiers were tested using the un-scrambled, correctly annotated, test sets. The green circles represent the performance of the classifiers when trained using un-scrambled annotations. Errors represent ± SEM (*note*: error bars are present but not discernible in the graph).

## Discussion

Open-source machine learning tools for behavioral neuroscience are being released at an intensifying pace^31^. We have identified, and attempted to solve, several issues that prevent a general adoption of machine learning techniques within the field. In many scenarios the required hardware, programming and computational familiarity, and the sensitivity of methods to the experimental conditions, may result in experimenters persisting with manual annotation methods. To advance access to machine learning methods we built SimBA, a software interface that can be used on standard computers by individuals with limited coding experience to implement and generate supervised machine learning algorithms for complex social behavior. We have conscientiously selected a supervised approach, and incorporated a number of explainability and validation checkpoints^38,39^, to ensure that researchers unfamiliar with these approaches are confident in the accuracy and validity of the classifiers they generate. We have supplied <u>extensive documentation</u> and <u>use-case scenarios</u> to simplify the iterative process of tuning supervised predictive classifiers to researcher expectations.

Here we use SimBA to create a battery of 28 predictive classifiers for behavioral repertoires relevant for studying social motivations in rats and mice^62,63^. We make SimBA, together with the generated classifiers and detailed tutorials, available online to expedite implementation of machine scoring complex behaviors within labs that have limited prior programming experience. Social classifications are readily consolidated with the analysis of calcium indicators or electrophysiological responses and we aim to integrate such alignment options for prevalent neural recording systems within the SimBA graphical interface (Fig. 11).

**Figure 11.**
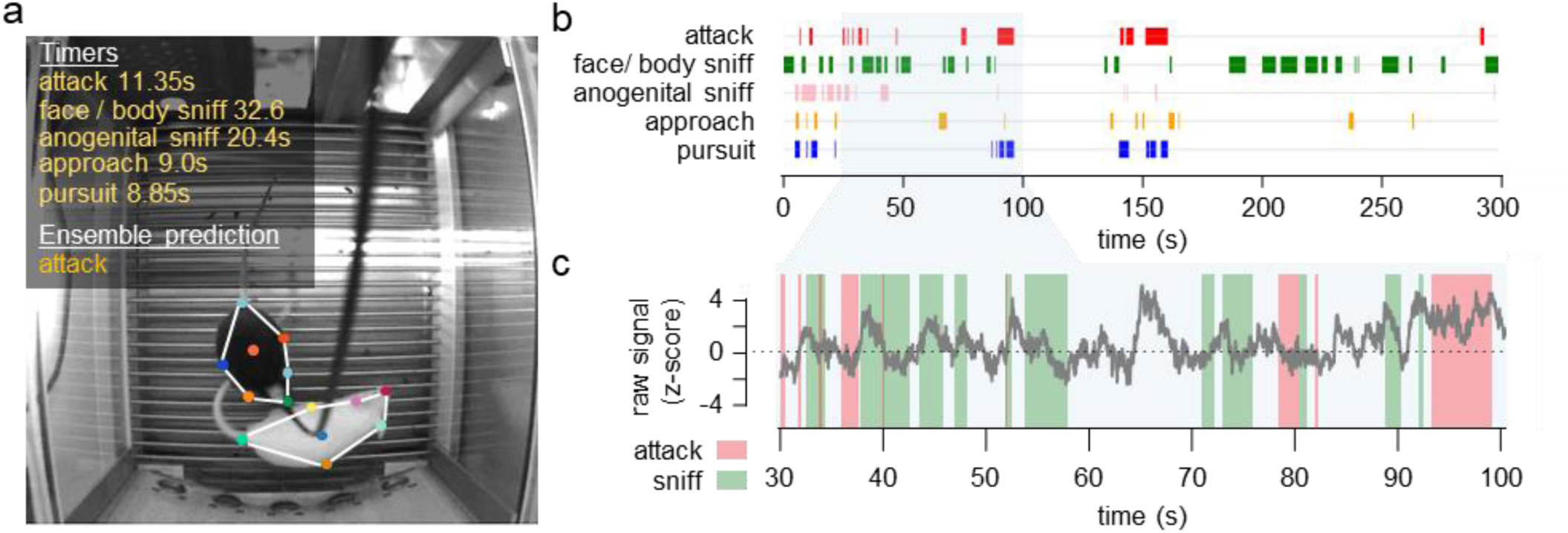
SimBA-generated social predictive classifications aligned with nucleus accumbens (NAc) fiber photometry traces. (a) The mouse resident-intruder protocol within an operant chamber, using a black CD-1 x C57 hybrid resident mouse expressing GRAB_DA_ in the NAc and a white BALB/c intruder mouse. (b) Gantt-plots created in SimBA that display the duration and frequency of classified social behaviors. (c) Z-scored fiber photometry traces (500-550nm) within a 70s-time window. Inset color bars represent classified bouts of attacks (red) and face / body sniffs (green).

We make the methods accessible by incorporating video pre-processing options, software support for importing pre-defined machine learning hyperparameters, performing common model evaluation techniques, and tools for visualizing the machine learning classifications. We demonstrate that SimBA is flexible by generating social predictive classifiers across multiple labs for both mice and rats, and by creating accurate classifiers from a third-party dataset available online^19,47^.

We used SimBA to create classification performance learning curves, where different percentages of the total behavioral annotations are used to create the random forest classifiers (Figs. 5–7). The learning curves show that, for most classifiers, significant improvements can be achieved by adding further experimenter-made behavioral annotations. Many labs working with social behavior already possess video data with accompanying annotations, and this data can be used to both expand the battery of classifiers and sharpen the performance of the classifiers generated here. We have made the datasets used to create our current classifiers available online, and we hope that others will consider further data sharing efforts that increase the scope and accuracy of social machine classifications.

Classifier performance improvements can also be achieved through statistical techniques. Accurate prediction of behavior requires handling class imbalances^64^. Datasets in biomedical and behavioral sciences will, in nearly all scenarios, be imbalanced such that observations of the majority class (e.g., observations of the absence of the behavior of interest) will substantially outweigh observations of the minority class (e.g., observations of the presence of the behavior of interest). In our datasets, rare events introduce class imbalances of approximately 1:85 and it is reasonable to assume that alternative behaviors present similar or higher imbalance ratios. SimBA includes a variety of statistical re-balancing tools that address class imbalances^65–68^. The parameters of these rebalancing tools can be user-defined or imported from similar and previously successfully generated classifiers. Balancing parameters for all 28 classifiers are shown in Table 1, as representative starting points.

The classifiers generated for this manuscript use the DeepLabCut^27^ default network architectures for poseestimation positional data, which the subsequent machine learning features are derived from. There are, however, many other effective open-source solutions for animal tracking and pose-estimation^20,22,69^ and SimBA is agnostic to the tools used to extrapolate positional coordinates. SimBA also has an interface for DeepPoseKit^26^ that permits a range of alternative neural network architectures for pose-estimation that may offer speed and accuracy advantages^70^. Others have also successfully implemented YOLO-based approaches^71^. Forthcoming releases of tracking packages promises individual identification of similarly coat-colored animals interacting in groups and we look forward to incorporating such significant developments into SimBA as they are released.

Social interactions - even between differently coat-coloured animals – introduce tracking and pose-estimation challenges that become aggregated by the turbulent interactions and substantial occlusion present in social behavior protocols. For example, DeepLabCut and DeepPoseKit occasionally attribute body-parts to the incorrect animal despite extensive annotation data. We approached the issue post-hoc and designed body-part correction tools that identify and correct movements and distances that are unrealistic based on experimenter-defined criteria. Although the two correction tools eliminate unambiguous outliers, further performance improvements may be obtained by more refined and flexible machine learning methods^72–74^ and/or data cleaning techniques that incorporates variable criteria for the different body-parts. We are currently working on evaluating such techniques and their potential benefits for generating random forest and other decision ensembles for rodent social behavior.

### Future directions and challenges

There are some important caveats to be aware of, or account for, during the creation of predictive classifiers with SimBA. Although SimBA can analyze two animals of different coat colors, or a single animal of any coat color, our method does not currently transfer to two or more animals of the same coat color. Current gold-standard methods for performing individual identifications in groups of similarly coat-colored individuals use RFID-data and depth camera systems^21,22^. Additional advances in machine vision has also addressed identification issues through dual segmentation/identification networks^75,76^ or by including measures of optical flow^77^ and part affinity fields^73^. Accurate mask-RCNNs^78^ and YOLO-based approaches^71^ may also be promising platforms for calculating the rich and relevant feature space that is required for individual identification in social behavior protocols. Identification issues could also be circumvented by adjusting experimental protocols and using differently coat colored mice of the same strain when possible^50^.

SimBA uses comprehensive and generic feature sets (e.g., movements and distances) in random forest designs to permit accurate classifications of variable user-defined target animal behaviors while avoiding critical multicollinearity issues^79^. In some use-cases, however, random forest designs may not be optimal, and nonspecialized feature sets may introduce decision ensemble noise which can negatively affect runtime and the interpretability of feature importance calculations. Commercial alternatives for generating predictive classifiers^80–82^ typically offer exhaustive automated feature engineering/selection methods and evaluate or combine larger batteries of classification techniques when searching for the best alternative. Such tools are also accessible through open-source packages^83–86^ and should be available in future SimBA releases to expedite processing, decrease algorithm complexity, and increase the flexibility of the toolkit. We have nevertheless found random forest designs to be the most robust for classifying the most typically scored aspects of aggression behavior.

A limit to accessibility is that body-part tracking and pose-estimation requires GPU processing. Cloud-based GPU solutions, such as Google Colaboratory / Microsoft Azure / Amazon Web Services, are promoted as alternatives to dedicated GPUs, but this can still be a challenging process. Providing pre-trained network weights only goes so far in addressing accessibility as users typically must add images representing their environment to the original neural network weights, as well as analyze their own new experimental videos. It is feasible that less complicated classifiers, such as with attack predictions that predominately rely on speed and distance metrics, can be accurately predicted (potentially in real-time) by combination of advanced computational methods such as optical flow^77^ and pretrained YOLO models^53^.

## Conclusions

We demonstrate that SimBA flexibly and simply creates accurate supervised machine learning classifiers using feature sets engineered from open-source pose-estimation tracking data. We demonstrate that the approach is robust by creating social classifiers for both rats and mice across multiple labs, and further by creating classifiers from a previously released public datasets available online^19^. We built a GUI that makes our method accessible to experimenters regardless of previous programming experience. This is a step towards generating a standardized toolbox of computational models for scoring complex social behaviors in preclinical neuroscience.

## Notes

<u>Funding and disclosure:</u> The authors declare that they do not have any conflicts of interest (financial or otherwise) related to the text of the paper. The research was supported by NIDA 4R00DA045662-02 (SAG), NIDA P30 DA048736 (SRON and SAG), NARSAD Young Investigator Award 27082 (SAG), and NIDA T32 5T32NS099578-04 (NLG). We thank Briana Smith, Liana Bloom, Annette Mercedes, Roёl Vrooman and Kayla Pitts for their thoughtful assistance annotating behavioral frames and discussing the manuscript and online documentation.

https://github.com/sgoldenlab/simba

https://osf.io/tmu6y/

### Competing Interest Statement

The authors have declared no competing interest.

